# Controlling membrane barrier during bacterial type-III protein secretion

**DOI:** 10.1101/2020.11.25.397760

**Authors:** Svenja Hüsing, Ulf van Look, Alina Guse, Eric J. C. Gálvez, Emmanuelle Charpentier, David F. Blair, Marc Erhardt, Thibaud Renault

## Abstract

Type-III secretion systems (T3SSs) of the bacterial flagellum and the evolutionarily related injectisome are capable of translocating proteins with a remarkable speed of several thousand amino acids per second. Here, we investigated how T3SSs are able to transport proteins at such a high rate while preventing the leakage of small molecules. Our mutational and evolutionary analyses demonstrate that an ensemble of conserved methionine residues at the cytoplasmic side of the T3SS channel create a deformable gasket (M-gasket) around fast-moving substrates undergoing export. The unique physicochemical features of the M-gasket are crucial to preserve the membrane barrier, to accommodate local conformational changes during active secretion, and to maintain stability of the secretion pore in cooperation with a plug domain (R-plug) and a network of salt-bridges. The conservation of the M-gasket, R-plug, and salt-bridge network suggests a universal mechanism by which the membrane integrity is maintained during high-speed protein translocation in all T3SSs.

## Introduction

Bacterial type-III secretion is a conserved mechanism that enables assembly of two large nanomachines in the bacterial cell envelope. The bacterial flagellum (assembled by the flagellar type-III secretion system, fT3SS) is involved in cellular motility through the rotation of a long extracellular filament. Injectisomes (assembled by the virulent T3SS, vT3SS) are present only in pathogenic Gram-negative bacteria and are used to deliver effector proteins into their eukaryotic hosts. We recently demonstrated that secretion of flagellin via the fT3SS of *Salmonella enterica* occurs through an injection-diffusion mechanism with a remarkably fast initial injection speed – up to tens of thousands amino acids per second – (1), and similar results were reported in *Vibrio alginolyticus* (2). This extreme secretion speed, without equivalent in other pore-based protein channels, raises the question of how membrane gating is preserved. In other words, how does the T3SS secretion pore enable high-speed translocation of proteins but not of water, ions and small molecules, to preserve membrane gradients and cellular physiology?

Bacteria have evolved several strategies to gate transmembrane channels and transporters. One common feature is the presence of specialized amino acids in the channel of the transporter. For instance, gating in the bacterial Sec translocon is warranted by a stretch of conserved isoleucines in the central constriction of the SecY channel (3), by four glutamines in the MacAB-TolC macrolide efflux pump and eight tyrosines in the Wza polysaccharide transporter (4). Another approach is the acquisition of gating domains, such as the outward facing plug domain of the SecY translocon (3). The core of the fT3SS export apparatus (EA) is composed of three proteins, FliP, FliQ, and FliR (SpaP/SctR, SpaQ/SctS, SpaR/SctT in the homologous vT3SS), which form a pseudohelical secretion pore complex. We previously reported that mutations in a conserved loop of methionines in FliP (M-loop) cause leakage of ions and small molecules through the membrane (5). However, the mechanism of secretion pore gating and the potential contribution of other components of the core EA remained elusive. In this work, we performed extensive mutagenesis and functional analyses to systematically investigate the role of FliP/Q/R and of conserved protein features of the secretion pore in the maintenance of the membrane barrier during type-III secretion.

Our results demonstrate that a plug domain in FliR (R-plug) and the M-loop in FliP cooperate to gate the fully assembled T3SS secretion pore. We further show that the ensemble of M-loops of all FliP subunits in the assembled secretion pore function as a deformable gasket (M-gasket), and that the stability of the secretion pore is maintained by a network of salt bridges in FliQ. The composition of the residues forming the M-gasket required the unique combination of physicochemical features that is found in methionines in order to support both the gating function and the conformational changes that are required for protein substrate secretion. Finally, we demonstrate that mutations compromising the membrane barrier during type-III secretion are detrimental to bacterial fitness.

## Results

### The T3SS secretion pore harbours several domains that contribute to seal the secretion channel

We previously demonstrated that FliP alone can form a pore under over-expression conditions (5). Accordingly, purified FliP oligomers displayed a decreased central density by electron microscopy reminiscent of a pore (6, 7) (Figure S1A). Recent structural studies have revealed that the functional EA is actually a hetero-oligomer, composed of FliP/Q/R with a 5:4:1 stoichiometry, and forms a helical secretion pore complex (8) (Figure S1B). An interesting feature of the T3SS pore is its organization in a two-level hetero-oligomer with a pseudo hexameric symmetry. The periplasm-facing top level is composed of five FliP subunits and of the N-terminus of FliR, while the bottom level is composed of four FliQ subunits, the C-terminus of FliR and one FliP subunit (P5) that is common to both levels (Figure 1A). This peculiar organization is enabled by a striking structural homology in two regions of FliP and FliR (residues 64–100/18–55 and 173–245/71–155), and between FliQ and the C-terminal part of FliR (residues 167–255) (Figure 1C). Interestingly, the strong structural homology between FliP and FliR is disrupted in the region that faces the central channel, between two α-helices. While FliP features a short, highly conserved loop of three methionines (residues 209–211) – for which we previously reported evidence that it acts as a gasket to close the T3SS channel (5) – FliR exposes instead a bulky domain (residues 106–120) that was hypothesized to act as a plug to seal the channel (8) (Figure 1B–C).

**Figure 1.**
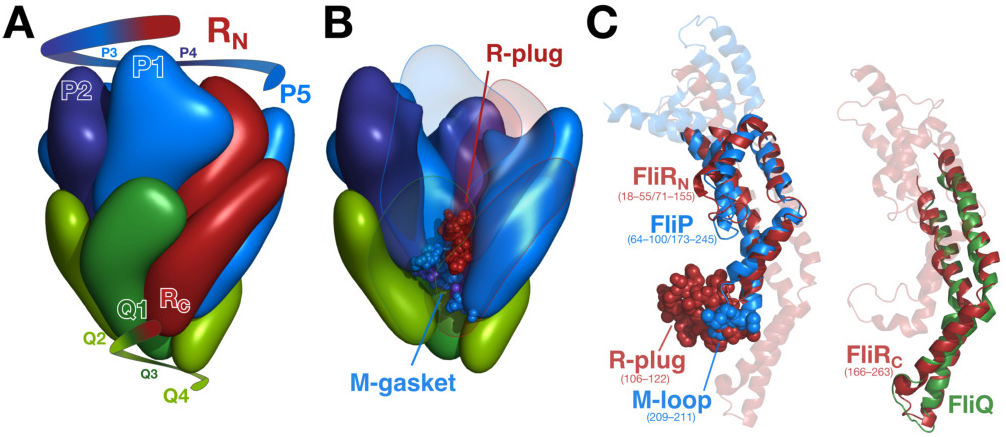
Organization of the fT3SS pore. **A.** The export gate is composed of FliP/Q/R in 5:4:1 stoichiometry and organized in a corkscrew-like helical structure. The R subunit has structural homology that is split between P (R_N_) and Q (R_C_) [see panel C].. **B.** The FliR plug (R-plug) and the M-gasket formed of 5 FliP M-loops are located in the core of the pore, sealing it in the closed conformation. **C.** The N-terminal part of FliR exhibits a strong structural homology with FliP, with the exception of a bulky domain (residues 106–122) in place of the FliP M-gasket. The C-terminus of FliR is similar to FliQ on its full length. The structures in A and B are based on the 6f2d PDB entry. The isosurface at downscaled resolution (8Å) was computed to to better illustrate the global shape and organization of the subunits.

Our previous work and structural data from others suggest that the T3SS EA has evolved specialized domains to preserve the membrane barrier. Accordingly, we first investigated the individual roles of FliP, FliQ, and FliR, and their cooperation in maintaining the membrane barrier during assembly and substrate export via the fT3SS using a combination of mutagenesis approaches and functional assays (Figure S1C).

### A highly conserved methionine loop in FliP plays a dual role to support channel opening while gating the pore

#### The methionine loop in FliP regulates membrane gating

We previously reported that FliP can form a pore in the inner membrane, both by itself and in combination with other flagellar proteins (now identified to be FliQ and FliR), and described a loop of three methionines (M-loop, residues 209–211 in *S. enterica* FliP) that appeared to play a role in membrane gating by forming an inter-molecular structure that we named the M-gasket. Deletion of a single methionine in the M-loop or mutagenesis of the triplet to alanines caused a drastic increase in membrane permeability (5). Our initial results suggested that the length of the amino acid side chain was important for the function of the M-gasket. Thereafter, several structural studies have confirmed that the M-loop is indeed exposed to the channel in both fT3SS and vT3SS (8–10). In the FliPQR secretion pore, five copies of FliP are present, which results in a local pool of 15 methionine residues that are spread over about 20Å at the base of the channel (Figure 2A). This high local enrichment in methionines and the strong conservation of the motif across distant bacterial species (Figure 2B) raises the question whether methionines have a specific property that could participate in the gating of the T3SS secretion pore.

**Figure 2.**
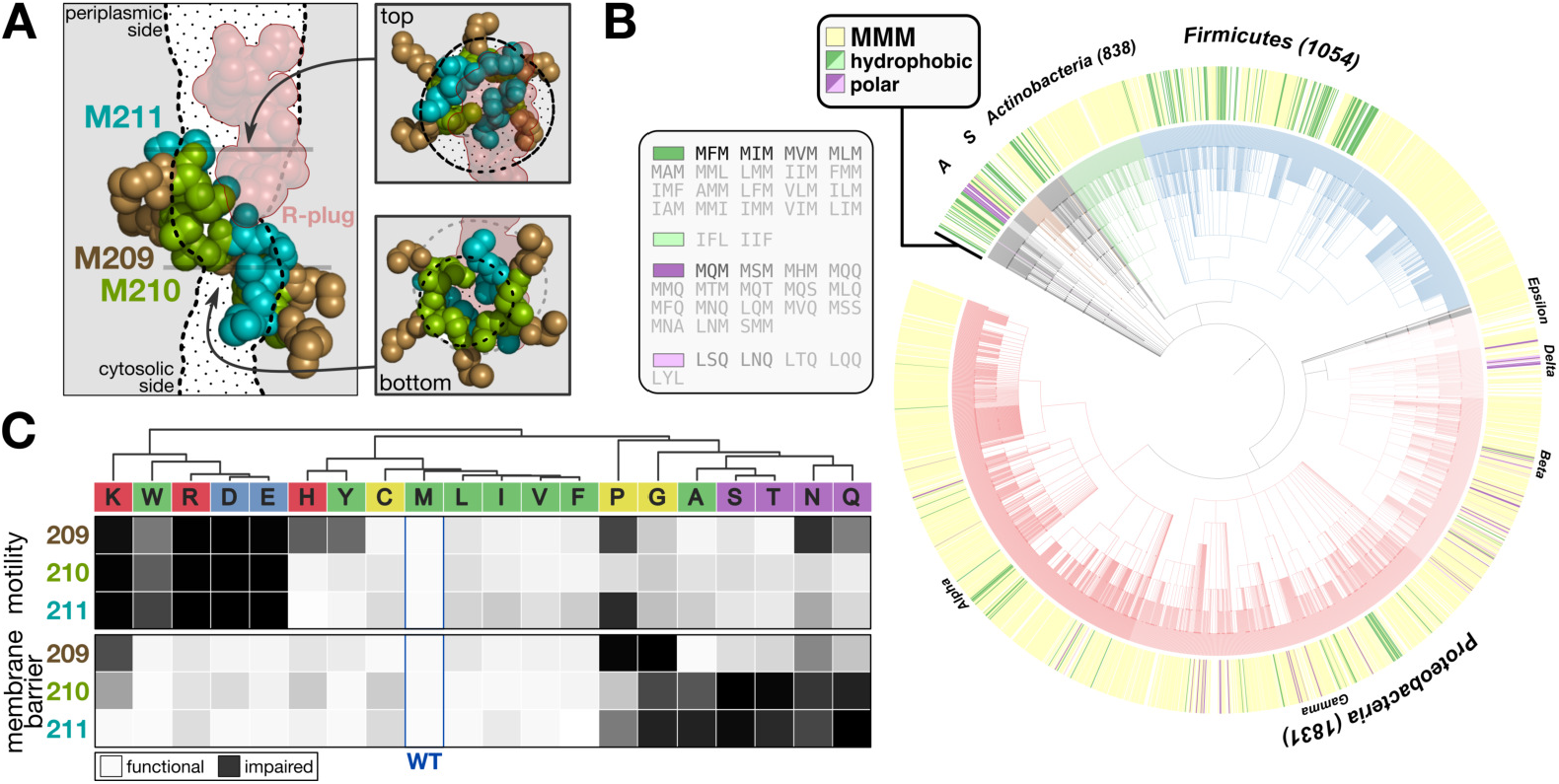
Conserved methionines in the PQR pore play a crucial role for T3SS function and gating of the pore. **A.** Positions of the conserved M-loops that form the M-gasket relative to the channel of PQR pore. The channel was computed using Caver 3 from the PDB:6f2d structure devoid of the M-loops and R-plug residues to model pore opening, and cross-sections were represented as dotted lines and surfaces. The methionine residues are color coded according to their position in the primary FliP sequence. **B.** Conservation of the M-loop in flagellated bacteria. The outer ring of the tree is colored according to the M-loop motif. The most conserved MMM motif is colored in yellow, other hydrophobic motifs in green, and polar motifs in purple (with a lighter color when the motif contains no methionine). The color in the central part of the tree represent the major represented phyla: Proteobacteria (decreasing shades of red corresponding to the alpha, gamma, beta, delta, and epsilon classes), Firmicutes (blue), Actinobacteria (green), Spirochaetes (S, brown), Acidobacteria (A, grey), and other phyla in black. The alternative motifs are listed on the left of the tree and shaded according to their frequency from black (MFM, 125 occurrences) to light grey (1 occurrence). **C.** Clustermap of the motility and membrane barrier function scores for all combinations of single amino acid substitutions at the three methionine positions. Amino acids are colored according to classical physicochemical groups (red: positively charged, blue: negatively charged, purple: polar, green: hydrophobic, yellow: special)

To unambiguously determine the biochemical properties required for this gating function, we performed a systematic replacement of each of the three methionine residues to all 19 other amino acids in FliP, which was expressed from its native chromosomal location. All mutant FliP proteins were expressed and found in the membrane fraction in levels similar to the WT, suggesting that single-amino acid substitutions did not affect FliP stability or complex formation in the inner membrane (Figure S2A). We next performed three complementary assays to determine the impact of the single amino acid mutations in the M-gasket on the function of the T3SS (Figure S1C): (i) motility in soft agar reported the final assembly of the flagellum, reflecting thus the overall assembly and secretion capability of the fT3SS; (ii) secretion of a fusion construct of the hook subunit (FlgE) to the TEM-1 β-lactamase (FlgE-bla) in order to quantify the ability of the T3SS to secrete an early flagellar substrate (11); and (iii) optical density changes in presence of high salt, reflecting the flux of water and ions across the T3SS secretion pore and thus reporting membrane barrier function (5).

Most single substitutions in the M-gasket allowed flagellar assembly and bacterial motility (Figure S2B–D). Strictly charged residues were not tolerated at any position, almost fully abrogating motility. Substitutions to histidine, however, only affected motility on the first methionine position, probably as histidines are mostly unprotonated at cytosolic pH thus greatly reducing their hydrophilicity. Interestingly, a genetic screen for pseudo-revertants led to the isolation of one fairly motile suppressor mutant (M-loop motif MER) bearing a combination of a positive and a negative charge (Figure S2E; Supplementary Table S1). Substitutions to large polar residues (histidine, asparagine, glutamine, and tyrosine) on the first position, rigid prolines on the first and third positions, and bulky tryptophans at all positions were only weakly tolerated. In this study we used motility as a proxy of substrate secretion over the whole set of fT3SS substrates as it is a simple and reproducible method. To confirm our approach we also monitored translocation of the hook protein FlgE into the periplasm and obtained similar results (Figure S2F).

In parallel, we tested membrane permeability in presence of a high concentration of guanidinium (0.5 M). Ion leakage was reported by the immediate change in optical density (OD_600_). This showed us that preserving the membrane barrier function by the fT3SS pore was mostly dependent on the second and third positions (the residues most exposed in the channel) and that leakage was promoted by substitutions to polar (S, T, N, Q) and small (A, G) residues, and by proline (P) (Figure S2D).

The motility and permeability data were combined and clustered by amino acid (Figure 2C). Three clusters were obtained which revealed the relationship between the amino acids properties and function of the T3SS. The first cluster regrouped strictly charged amino acids (R, K, D, E) and bulky tryptophan (W). Substitution to those residues abrogated secretion and motility but had limited impact on membrane permeability, suggesting that the mutants could either be impaired for pore assembly or stability, or prevent translocation of the substrates. The second cluster comprised the polar uncharged amino acids (S, T, N, Q) and the small alanine and glycine residues (A, G). Mutation to those residues moderately affected motility and secretion but strongly increased permeability to small ions through the membrane. This suggested proper assembly of the pore, allowing substrate secretion, but an impaired ability to regulate gating of the pore. Finally, a third group, clustering the hydrophobic medium to large-sized amino acids (V, I, L, M, F, Y), cysteine (C) and histidine (H), had a rather neutral effect on both parameters. Overall, loss-of-function mutations on the first position (209) affected secretion and motility, but only moderately affected the maintenance of the membrane barrier. This suggests a structural role of this residue, which is well supported by the structural data. In the structure of the closed pore reported by Kuhlen *et al.*, the residue is indeed buried in the PQR complex (8). Conversely, the residues at the second (210) and third (211) positions are directly exposed to the channel. Apart from mutations to charged residues, and structure-destabilizing residues (proline, glycine, and tryptophan), substitutions on positions 210 and 211 had a rather limited impact on secretion and motility, but the presence of polar residues strongly reduced the ability of the pore to gate small ions. Hydrophobic residues of medium to large size were well tolerated at all positions, indicating that hydrophobicity of this domain is an important factor to keep the pore sealed.

#### The physicochemical features of methionine are required for the M-gasket to function as a deformable gasket

Our single substitution approach indicated that hydrophobic amino acids of medium size side-chain length were required in the pore-exposed M-gasket of FliP in order for the assembled pore to facilitate both secretion of substrate proteins and preserving the membrane barrier. Interestingly, despite a very strong conservation (Figure 3A), our results further suggested that methionines are not strictly required at any particular position in the M-loop (Figure 2B). However, in the context of single substitutions, two methionines are still present per subunit for one mutated residue. Due to the helical nature of the channel, each residue in the five M-loops has a slightly different position in the M-gasket and different neighbors. Furthermore, while the structure of the open pore remains currently unknown, it is obvious that conformational changes are required to allow substrate translocation. Thus it appears reasonable to assume that the global – rather than position specific – residue composition is important.

**Figure 3.**
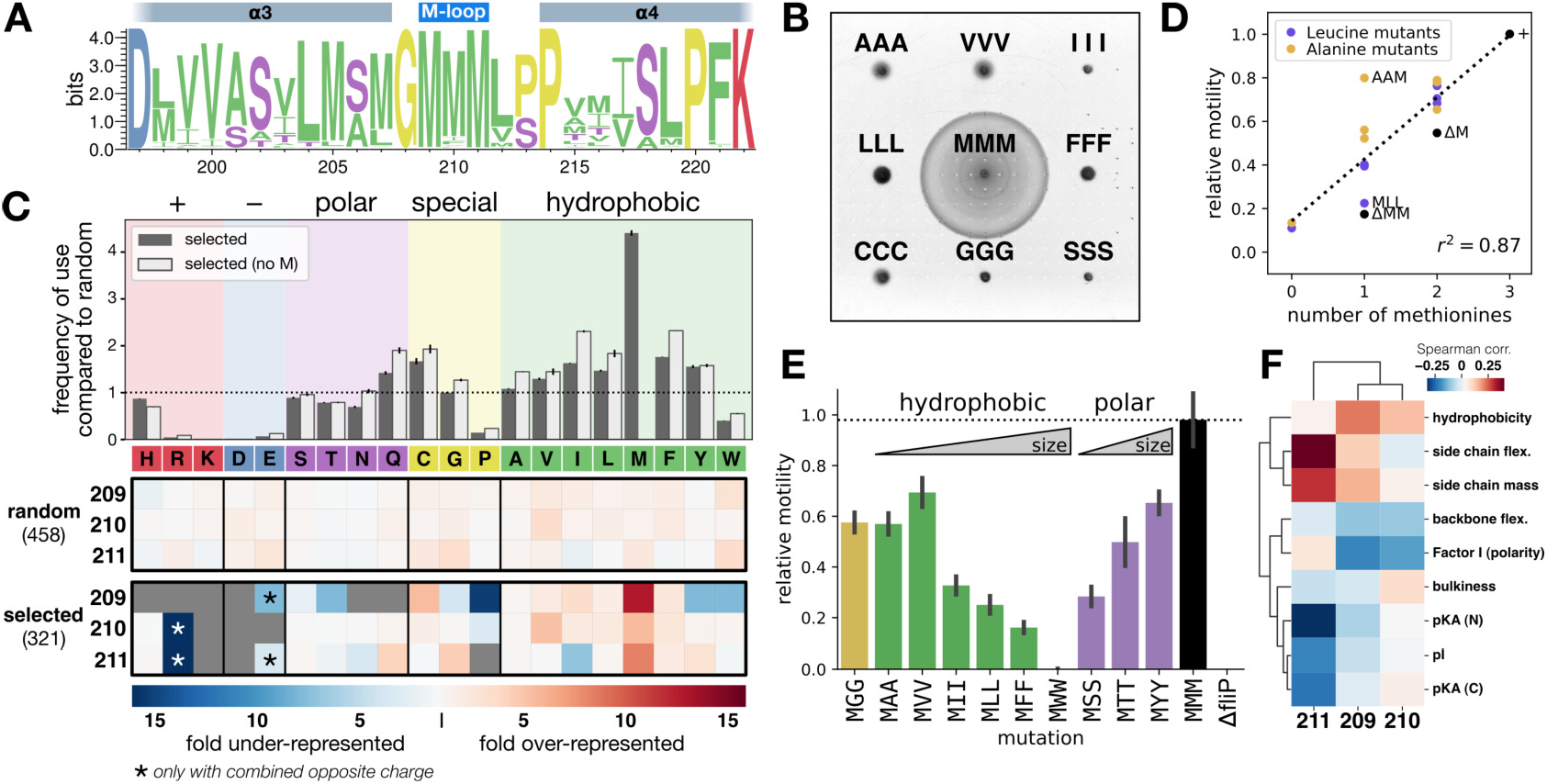
Methionine residues display a unique combination of physicochemical features that are required for T3SS function. **A.** Unlike the FliR plug, methionine residues in the FliP M-loop are highly conserved. From a selection of about 2,500 unique sequences of FliP homologs reported in a previous work (6), the first and last methionine residues are 99% conserved and the central one 87%. **B.** While single amino acid substitutions to small and hydrophobic residues have a limited impact on motility (Figure 2B), triple substitutions almost completely abrogate fT3SS function. **C.** When selected for fT3SS function, M-loop variants to random triplets display a strong bias towards specific residues. Hydrophobic residues are primarily selected, with a net preference for methionines. Position-specific variations highlight different contributions of the three residues to FliP function (see main text). Top: amino acid frequency of use in the M-loop for all residues. Methionines are strongly over-represented (dark grey bars). When only motifs devoid of methionines are considered (light grey bars), only a small increase of use is observed for medium size hydrophobic amino acids without a clear candidate to substitute for methionine residues. **D.** Overall, fT3SS function is proportional to the number of methionines in the motif (see Figure S3E). **E.** Side chain size of the exposed residues 210 and 211 impacts fT3SS function differently depending on the nature of the amino acids. Large hydrophobic residues and small polar residues decrease fT3SS function. This suggests that function of the M-loop requires a non-linear, complex, combination of several physicochemical features. **F.** Position-specific correlation between fT3SS function (motility) and physicochemical features reveals that side-chain flexibility, amino acid-size, and hydrophobicity are key favorable features required in the M-loop for fT3SS function. In contrast, bulkiness, side chain polarity, and backbone flexibility are on average unfavorable. Spearman correlation was computed here, but similar results are observed using Pearson correlation or distance metrics. Experiments were performed with at least 3 biological replicates. Error bars represent the standard deviation of the calculated probability in panel C and the standard deviation of the mean in panel E.

In line with this hypothesis, we observed that triple sub-stitutions to hydrophobic amino acids other than methio-nine almost fully abrogated flagellar assembly and motility, suggesting a special role of methionine stereochemistry in this domain, which is not restricted to hydrophobicity (Figure 3B). Single position mutagenesis had allowed us to identify biochemical properties that were causing defects in T3SS secretion and gating functions when the rest of the M-loop remained unchanged (i.e., one substitution per two methionines). To resolve the properties that are required for those functions in a general context, we generated random chromosomal variants of the M-loop by PCR using degenerate oligonucleotides. This differed from the single-position mutagenesis experiment in the fact that, in this approach, individual amino acid contributions were challenged in a random, non-optimized context. This setup is thus more likely to reveal important physicochemical requirements of residues forming the gasket of the secretion pore.

To obtain full amino acid coverage while minimizing genetic code redundancy, rare and stop codons, we used a (NNS)×3 mutagenic sequence of the M-loop. The variants had arbitrary amino acid triplets in place of the triplet of methionines, and were either isolated without selection (i.e. allowing any arbitrary combination), or selected to retain some degree of T3SS function using three different approaches (Figure 3C; Figure S3A–B). As expected, the non-selected variants revealed a homogeneous distribution of all amino acids, matching the frequencies of the NNS codons (Figure 3C). On the contrary, when we selected for combinations of M-loop residues that retained fT3SS function, we observed a clear bias in the frequency of certain residues (Figure 3C). Interestingly, the number of functional variants was much lower than the frequency that is expected considering that a functional motif could be any combination of amino acids that did not strongly affect motility (Figure 2B and Figure S3C). This indicated that the actual combination of residues in the motif is an important parameter to assemble a functional and gated secretion pore.

At first glance, the pattern was very similar to that obtained with the single substitutions: for example, charged amino acids were virtually absent; polar amino acids, proline, glycine, and tryptophan were obtained with lower frequencies, in particular on the first position. There were, however striking differences compared to the single substitution dataset. For instance, in few occasions charged residues were well tolerated, leading to motility close to that of the WT (Figure S3D), but only when the opposite charge was also present in the motif (e.g., EGR, CRE, AHE). Those combinations likely result in a null net charge, which confirms that a net charge is detrimental for fT3SS function, as observed in the single substitution dataset. Stringency towards specific amino acids was also substantially more apparent than in the single substitutions experiments. As an example, only medium size hydrophobic amino acids were permissive on the first position, but not large or polar ones. Most importantly, methionines were clearly over-represented in the motifs of the selected variants, which validated that stereochemistry of methionines has unique features required for the proper function of the M-gasket. (Figure 3C).

To assess the influence of the number of methionines on fT3SS function, we generated variants with all combinations of methionines with a given other amino acid. We chose alanine and leucine as those two hydrophobic residues demonstrated rather neutral changes in frequency in our random variants, but have different sizes. Furthermore, mutation from methionine to leucine is the most commonly found natural variation in proteins. Overall T3SS function was proportional to the number of methionines in the motif (Figure 3D + Figure S3E). When a single methionine was present in the motif, alanine constituted a better partner than leucine. This emphasized the importance of side chain size and flexibility of the motif, presumably to accommodate channel conformational changes when substrates are traveling through. We thus generated mutants of the two residues that are exposed in the channel (M210, M211) to hydrophobic and polar residues of various side chain size (Figure 3E). Among substitutions to hydrophobic amino acids, increasing the size of the side chain proportionally reduced motility, yet the opposite effect was observed for polar substitutions. This indicated that while side-chain size is an important factor, it is also context dependent. The most illustrative example is perhaps the difference of function between the MFF and MYY mutants. Phenylalanine, a large hydrophobic residue, resulted in a weakly motile phenotype, while tyrosine – which possesses an extra hydroxyl group, and thus has an amphipathic character – resulted in approximately 70% motility compared to the WT.

Taken together our data highlight that a non linear, complex combination of physicochemical features is required in this channel-exposed M-gasket to support proper fT3SS function. In an attempt to identify those features in an unbiased manner, we next assessed the function of several hundred unique motifs among our random variants (Figure 3C) using a medium throughput motility assay (Figure S1C). In line with our other data, correlation between motility and various physicochemical features indicated that side-chain flexibility, intermediate size, and moderate hydrophobicity are important factors for fT3SS function (Figure 3F).

We therefore propose that methionine residues combine ideal physicochemical properties in order to create a deformable gasket that accommodates the structural constraints of the T3SS pore, and allows high-speed translocation of substrates, while preventing leakage of small molecules. Interestingly, flexibility of the methionine side chain had been identified as a key property in the eukaryotic calmodulin, enabling local structural changes to accommodate binding to a large variety of targets (12).

#### The FliR plug and FliP M-gasket cooperate to seal the T3SS pore

The plug of FliR (R-plug) was identified in the previously reported FliPQR structure as a domain composed of large hydrophobic residues and proposed to act as a gating domain as it occludes the channel in its closed conformation (8), but it remained to be functionally characterized. We therefore performed directed mutagenesis of the plug domain, in order to determine the potential role of the R plug in gating of the T3SS channel. We generated several mutants that destabilize this domain by either substituting the large or rigid amino acids of the domain (residues 109–120, SFATFVDPGSHL) with smaller flexible residues (SAATAASGGSSA), or replacing the domain with short flexible serine/glycine loops (SGS, SGSGS, or SGSGSGS) (Figure 4A). All mutations resulted in a strong permeability to ions across the membrane (Figure 4B + Figure S4C). The plug mutant strains, however, did not demonstrate a defect in flagellar function (Figure 4C). Despite the loss of membrane barrier, the mutants displayed a motility phenotype similar to that of the WT, even in the presence of high concentrations of ions in the medium, suggesting that assembly and function of the fT3SS was not impaired (Figure S4D).

**Figure 4.**
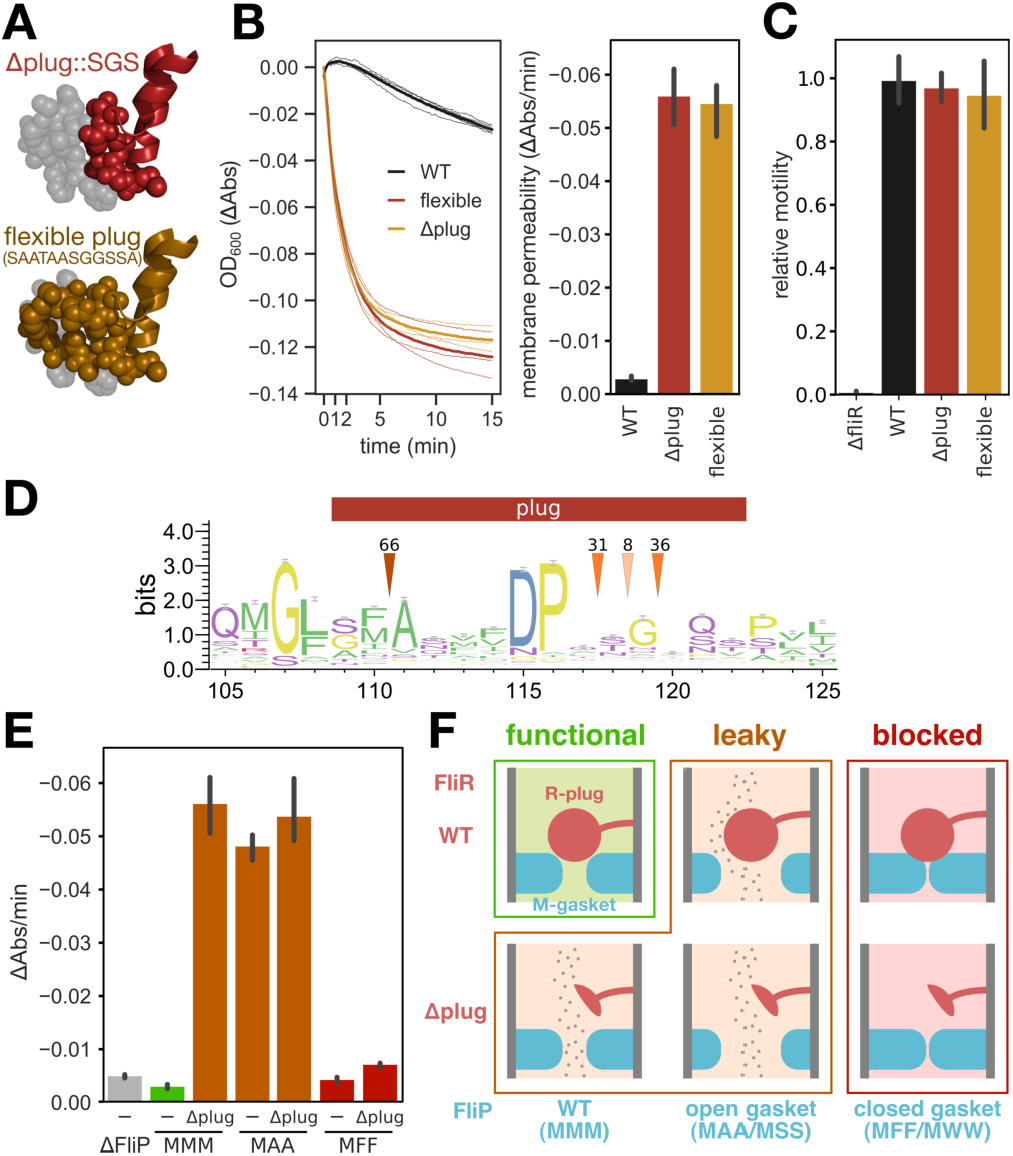
The FliR plug cooperates with the M-gasket to seal the T3SS pore and maintain membrane barrier. **A.** Mutations in the plug domain of FliR. replacement of residues 109–120 (SFATFVDPGSHL) with SGS (top) and substitution of the large amino acids to smaller flexible ones (SAATAASGGSSA, bottom). The WT plug is shown in grey for comparison (PDB: 6f2d). **B.** Both R-plug mutations induce a strong loss of membrane barrier as reported by the drop of OD_600_ in presence of a high salt concentration. **C.** Mutations of the plug domain do not impair flagellar assembly and motility. **D.** The plug domain is poorly conserved across FliR homologs (numbering according to FliR of *S.* Typhimurium). Arrows indicate the number of homologs with insertions at the pointed location. **E.** Single M-gasket and R-plug leaky mutations cause the same defect as the combination of both indicating that the two domains likely cooperate to gate the T3SS pore. Conversely, a mutation in the M-gasket that tightens the channel prevents leakage induced by a R-plug mutation. **F.** Model of the cooperation between the M-gasket and R-plug. Experiments were performed with at least 3 biological replicates. Error bars represent the standard deviation.

Accordingly, our data demonstrates that the plug domain of FliR plays an important role in gating of the EA pore. However, the absence of a direct impact on flagellar assembly and function suggests that tight membrane gating is not strictly required for protein translocation via the T3SS per se, but is rather important for the general physiology of the cell. Furthermore, this observation, the poor sequence conservation, and the high phylogenetic diversity (Figure 4D) supports the idea that the plug in FliR may have acquired a specialized role to seal the pore in its closed conformation.

The close proximity between the FliP M-gasket and the FliR plug (~3.5Å) raises the possibility that the two domains cooperate in gating of the channel (Figure S4A). If the M-gasket and the R-plug act as two independent gating domains, then it would be expected that a combination of both mutations is required to cause the greatest defect. In contrast, if the two domains act in cooperation, failure of either of them would be sufficient to impair the gating function. To address this issue, we measured the leakage of several combination of M-loop mutants in presence of a deletion of the R-plug.

The most leaky mutants of the M-loop and mutants of the FliR plug induced a comparable membrane permeability. We observed, however, no additive effect when both mutations were combined (Figure 4E–F and Figure S4C). This suggested that the two domains do not have a redundant function but rather are simultaneously required to gate the fT3SS pore, probably acting in cooperation. To further validate this observation, we introduced mutations to bulky amino acids in the M-loop to tighten the channel (MFF and MWW) and combined it with the deletion mutant of the FliR plug. Both M-loop mutations prevented leakage through the channel in the FliR plug mutant background (Figure 4E–F and Figure 4C).

### Control of membrane barrier is required at the level of the assembled and actively secreting complex

#### Intermediates of EA assembly are not leaky

The FliP/Q/R complex assembles sequentially. FliP oligomerizes first with the help of the integral membrane chaperone FliO, FliR is then recruited, and finally FliQ assembles at the bottom of the structure (6, 7). Deletion of a given component blocks assembly in one of the intermediate states (6). Our previous data indicate that FliP alone is able to form a stable pore when over-expressed from a plasmid, and that a ΔM mutation in the M-loop causes a detectable loss of membrane barrier function (5, 6). Yet, while this observation is interesting to decipher the steps of EA assembly, it does not necessarily reflect the regulation of the secretion pore under physiological expression conditions. We thus investigated whether EA assembly intermediates induced membrane permeability when expressed from their native chromosomal location and thus whether gating of the secretion pore played a role during the assembly of the T3SS.

For this, we introduced a leaky mutation (MAA) into the M-loop of various EA assembly intermediates. While the final complex was leaky in presence of the MAA mutation in FliP, none of the intermediates of assembly were sensitive to such mutations (Figure 5A). This indicated that the EA assembly intermediates are not able to form an open pore in the membrane. When combining the MAA variant with deletion mutants of flagellar proteins strictly required for T3SS function (FlhA, FlhB, FliF) or important for efficient substrate secretion (FliH/I/J ATPase complex (13, 14)), only mutations in the dispensable FliH/I/J ATPase complex caused leakage (Figure 5B). While the E211D mutation in FliI causes a much reduced motility, it allows to secrete many flagellar substrates to levels comparable to that of the WT (15) and interestingly, we measured that this mutation causes a leakage that is similar to that of the WT. This suggests that active substrate protein secretion is required for the T3SS secretion pore to open. Accordingly, another ATPase mutant (FliI E211Q), which was previously reported to display a much reduced secretion, demonstrated an intermediate leakage phenotype (Figure 5B). Our data thus suggest that only a completely assembled, and likely actively secreting, export apparatus is able to open its secretion pore, and that only the open state of the pore requires gating.

**Figure 5.**
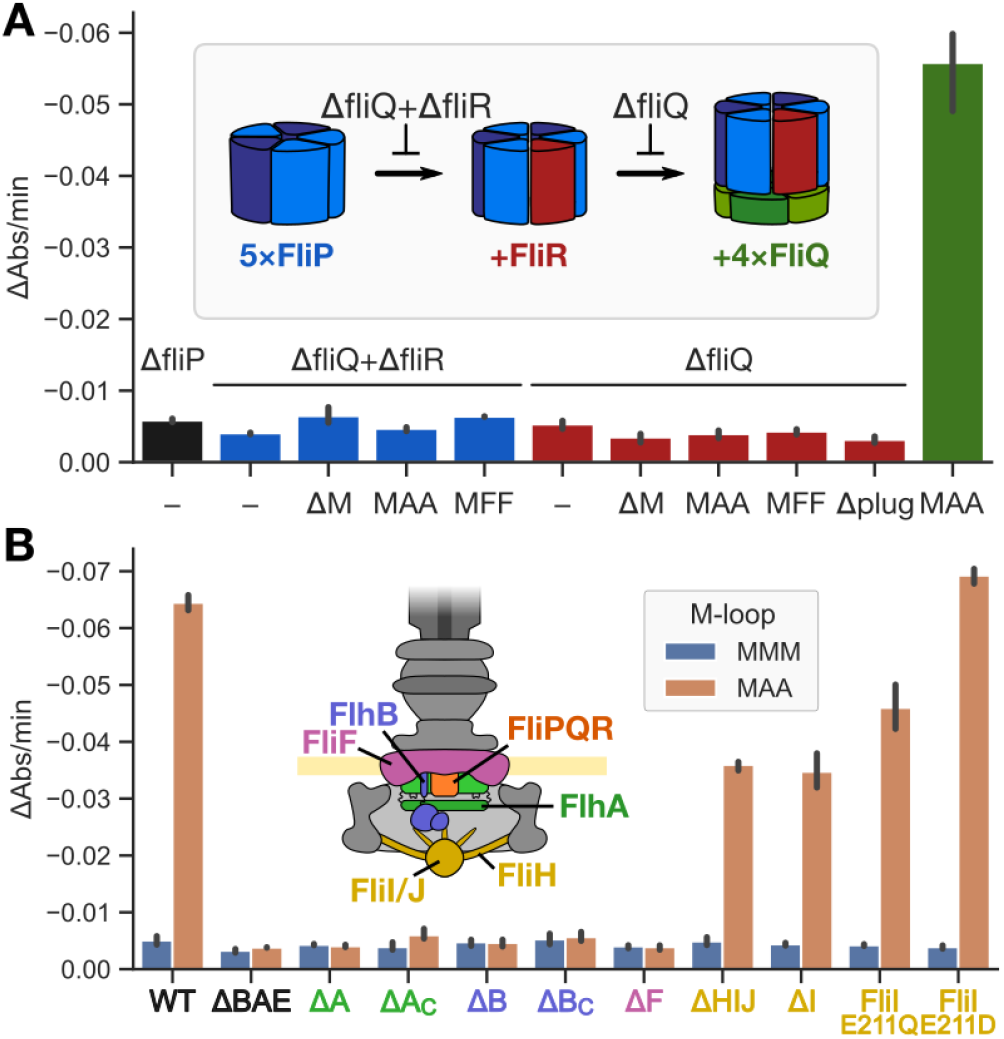
Intermediates of assembly do not need gating. **A.** Deletion of FliQ+R and of FliQ enables to lock the T3SS secretion pore in an intermediate assembly state (6). Using a leaky M-loop mutant (MAA) we determined that intermediates of secretion pore assembly cannot form an open pore and thus do need gating to preserve membrane barrier. **B.** Only functional T3SS secretion pores can open and need to be gated. The leaky MAA M-loop variant was combined with mutants of T3SS proteins. FlhA (A, A_C_: C-terminal domain), FlhB (B, B_C_: C-terminal domain), FliF (F) are all strictly required for T3SS function. FliH/I/J, that form the flagellar ATPase complex are however not strictly required. Only mutants of the ATPase complex, that retains some level of secretion, induce leakage in presence of the FliP MAA mutation. Experiments were performed with at least 3 biological replicates. Error bars represent the standard deviation.

### FliQ stabilizes the FliPQR complex

#### Systematic suppressor analysis of the M-loop reveals residues important for fT3SS secretion pore function and gating

Both the methionine loop in FliP and the plug domain in FliR are involved in gating of the fT3SS channel. While the plug in FliR appears to be dedicated to gating the channel, the role of the M-gasket is likely a trade-off between gating the channel and accommodating deformation of the gasket during secretion pore opening and substrate translocation. Our single substitutions experiment indicated that only the second and third methionines of the M-loop (M210, M211) are involved in gating the channel, while the first one (M209), which is buried in the protein structure, is rather involved in maintaining the stability of the complex (Figures 2B and 3C). Accordingly, the poorly functional AAA mutant assembles less flagella (Figure S5A) and is therefore less leaky than the single and double alanine mutants (Figure S3G). This idea was further reinforced by the analysis of motile revertants from the homogeneous triple mutants of the M-loop (Figure 6A–B). The almost non-motile triple M-loop mutants were inoculated in soft agar motility plates and incubated until flares of motile motility revertants were observed, which indicated the acquisition of a gain-of-function mutation. Interestingly, motile revertants of small residue M-loop mutants (GGG, AAA, SSS) were easily acquired and led exclusively to extragenic mutations in well characterized flagellar genetic regulators that are known to induce an over-expression of flagellar genes (Figure 6B, Supplementary Table S2). We hypothesize that over-expression of flagellar genes bypasses assembly defects of these triple M-loop mutants, similar to suppressor mutations bypassing the requirement of the fT3SS associated ATPase (16) or flagellar C-ring (17). Conversely, motile revertants of M-loop mutants consisting of larger and more rigid amino acids almost systematically reverted the initial M-loop mutation to incorporate smaller, more flexible or more polar residues, with the exception of the FFF variant for which we never isolated revertants (Figure 6B). Intriguingly, the GGG variant gave rise to a revertant bearing a FliQ E46D mutation. This gave us a first indication that FliQ could impact the function of the M-gasket.

**Figure 6.**
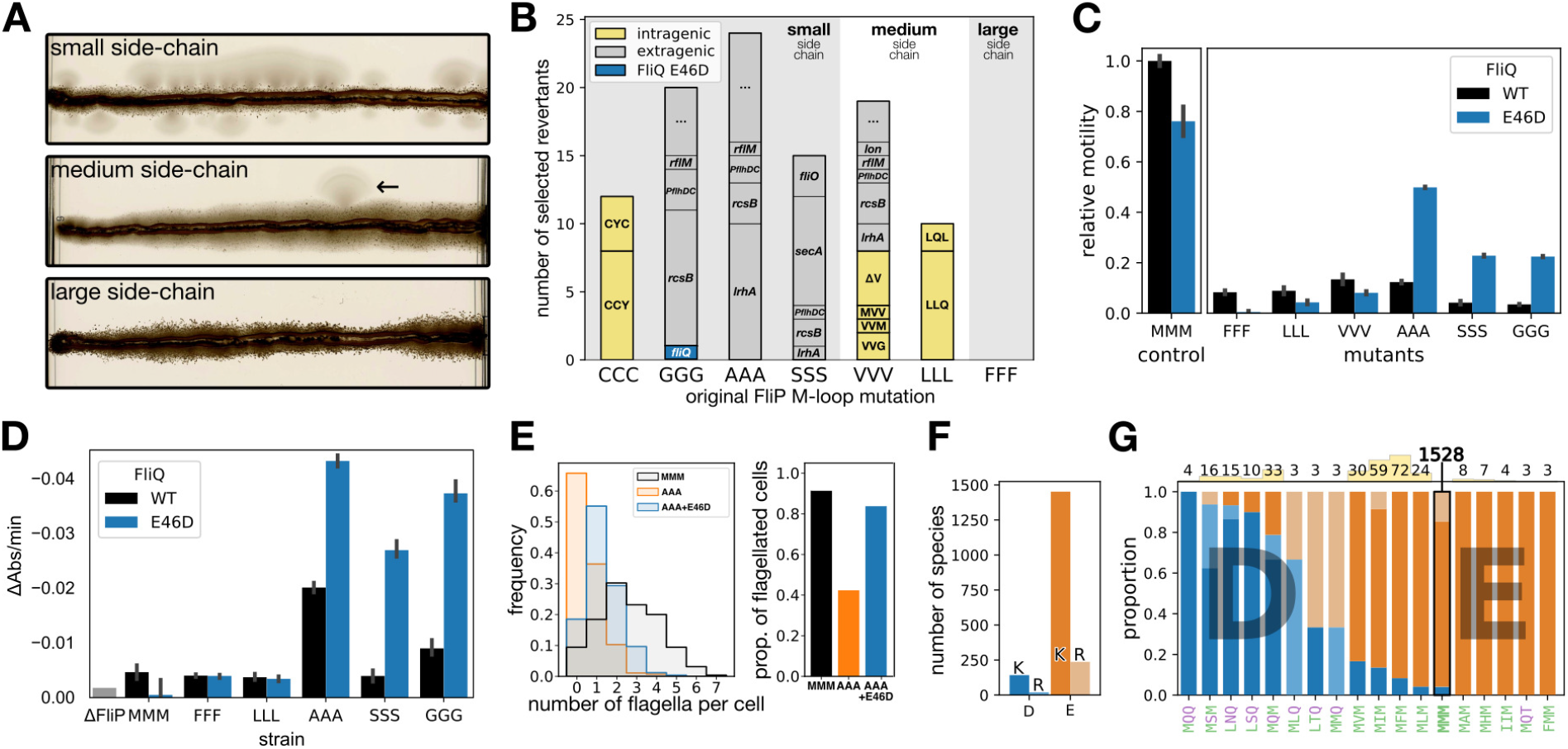
FliQ stabilizes the T3SS secretion pore through a network of salt bridges. **A.** Weakly motile mutants from Figure 3B were streaked into soft agar and monitored for several days until “flares” of motile revertants were obtained. M-loop mutants to small amino acids had a high reversion rate, medium size amino acids gave isolated revertants. We never obtained revertants for the mutant to large FFF residues. **B.** Small size M-loops revertants were almost systematically mutated in regulatory genes of flagellar assembly, with the exception of the CCC motif which we interpret as a possible channel constriction due to inter-cysteine di-sulfide bond formation. CCC and the medium size LLL required mutation of the M-loop to restore motility function. The intermediate size VVV acquired either extragenic mutations or M-loop reversions. Interestingly, M-loop mutations consisted in substitutions to amino acids with different physicochemical properties (polar vs hydrophobic, small vs large), acquisitions of methionines or deletion of one residue. This indicated that the CCC, VVV, and LLL mutants are likely reducing channel size, impairing substrate secretion, while the small, highly flexible GGG, AAA, and SSS motifs are probably leading to unstable protein complexes. Gain of function mutations identified in well-known regulators of flagellar assembly are represented in grey. Interestingly, the polar M-loop variant SSS gave rise to mutations in FliO and SecA, both of which are directly implicated in FliP assembly. “…” indicates that no obvious candidate was identified. **C.** The E46D mutation in FliQ has opposite effects depending on the background of the M-loop. Combined with the WT or with M-loop mutants with poorly flexible side chains, the E46D mutation decreases motility. In contrast, when combined to flexible M-loop variants that have an assembly defect, the E46D mutation improves motility. **D.** The stabilizing effect of the FliQ E46D mutation enhances leakage induced by the small and flexible M-loop mutants. **E.** The FliQ E46D mutation improves the flagellation of the defective AAA M-loop variant, both in terms of number of flagellar filaments per cell (left) and of proportion of flagellated bacteria (right). **F.** The majority of FliQ homologs use a glutamic acid (E) in their inter-molecular salt bridge combined with a lysine (K). **G.** The majority of FliQ homologs use a glutamic acid (E) in their inter-molecular salt bridge combined with a lysine (K). G. The small proportion of FliQ that have an aspartic acid (D) are strongly correlated with M-loop variants possessing a polar residue. The number of occurrences of each M-loop variant is indicated on the top of the graph and represented as a small yellow bar for comparison (except for the WT due to scale). Hydrophobic and polar residues of the M-loop are colored in green and purple, respectively. Salt bridges with lysine (K) and arginine (R) are colored in dark and light shade, respectively. Only motifs found in more than two occurrences have been included. Experiments were performed with at least 3 biological replicates. Error bars represent the standard deviation.

#### FliQ stabilizes the FliPQR complex

FliQ is located on the cytosolic side of the FliPQR pore. It assembles last into the FliPQR complex (6). While the protein is not required to form the initial, stable, FliPR complex, FliQ is essential to complete the assembly of the fT3SS secretion pore. FliQ possesses a relatively conserved methionine (M47) that is in close proximity to the FliP M-gasket (4.7±0.9Å). We initially hypothesized that this residue could contribute to the pool of methionines in the M-gasket and therefore participate in gating the pore. However, mutations were well tolerated (even to charged residues) and had only a limited impact on motility and on gating of the pore (Figure S5B–C). This suggested that the M47 was not required for M-gasket formation and fT3SS function but might rather participate in fine tuning of gating.

The four FliQ subunits form α-helical hairpins with a kink in their center. Their tight interaction is supported by inter-molecular salt bridges involving the residues E46 and K54, and it was hypothesized that they are important for the stability of the complex (8). Accordingly, we found that mutations of those residues to alanine almost fully impaired motility (Figure S5B), which confirmed that a tight interaction between the FliQ subunits is required for T3SS function.

Before the structure of the FliP/Q/R complex was known, we had performed tryptophan replacements in the two α-helices of FliQ to destabilize their structure and identify important residues and potential interfaces of interaction. We obtained two mutations that fully abrogated motility: A28W and L29W (Figure S5D), presumably by preventing EA assembly or substrate translocation. Interestingly, the L29W variant spontaneously gave rise to a motile revertant consisting of a single methionine deletion in the M-loop of FliP. This result was the second indication in our data of a possible interplay between FliQ subunit packing and the function of the M-gasket. We therefore suggest that FliQ might act as a clamp to maintain the stability of the secretion pore, and eventually indirectly contribute to gating of the secretion pore (Figure S5E). Given the small change resulting from the E to D mutation in the FliQ E46D mutant, we hypothesized that the smaller aspartic acid would pack the structure even further, and that this mutation should decrease fT3SS function in a WT background of the FliP M-loop. As expected, in this background the E46D mutation resulted in a slight decrease in motility (~20%) (Figure 6C).

To further assess the connection between FliQ packing and the function of the FliP M-gasket, we combined M-loop variants with the FliQ E46D mutation. Depending on the nature of the M-loop mutants, the FliQ E46D mutation had antagonistic effects. M-loop variants to large amino acids (FFF, LLL, VVV) behaved like WT FliP: motility was further decreased and no change in leakage was detected. Consistent with our hypothesis, the FliQ E46D mutation had the opposite effect in combination with M-loop mutants to small amino acids (GGG, AAA, SSS): fT3SS function was improved, and leakage was increased (Figure 6D, Figure S5F), which we attribute to enhanced stability of the PQR complex. Accordingly, the FliQ E46D mutation improved the flagellation of the AAA M-loop variant (Figure 6E, Figure S5A).

The FliQ residues involved in forming the inter-subunit salt bridges are conserved in all currently documented homologs. The vast majority utilize a glutamic acid (E), often in combination with a lysine (K), while the presence of an aspartic acid (D) remains rare (Figure 6F). Interestingly, we observed that the use of an aspartic acid correlates with the presence of hydrophobic residues in the M-gasket that are prone to induce ion leakage according to our data from *S.* Typhimurium (Figure 6G). The exact impact of the presence of hydrophobic residues in the M-loops of these species would need to be investigated, however the observed correlation suggests that the FliQ E46D variant might have co-evolved with the M-loop to maintain the membrane barrier for these M-loop compositions.

In summary, our data indicates that the M-gasket has a dual role in gating the T3SS secretion pore and accommodating conformational changes required for high-speed substrate translocation. The latter function likely requires a deformation of the structure of the M-gasket that challenges the stability of the secretion pore. We observed that FliQ packing impacts the bulk of the M-gasket (and thus, likely the size of the secretion pore) and, reciprocally, that instability triggered by flexible M-loops can be corrected by a reorganization of the inter-FliQ salt bridges network. This indicates that FliQ plays a crucial role in maintaining the stability of the T3SS pore, and indirectly in maintaining the membrane barrier.

#### Physiological role of T3SS gating

Our extensive functional analysis of conserved features of the T3SS export apparatus demonstrates that gating of the secretion pore is achieved by a combination of several factors: a flexible deformable M-gasket made of the M-loops of five FliP sub-units, a bulky plug domain in FliR that cooperates with the M-gasket to seal the pore, and the stabilization of the complex by a network of salt bridges in FliQ. Furthermore, our results indicate that the export gate is likely a dynamic structure that supports conformational changes during secretion of substrate proteins (Figure 7A). Our analysis of FliR plug and FliP M-loop mutants demonstrated that those two domains prevent passage of water and ions across the secretion pore. Loss of membrane barrier results in alteration of bacterial physiology through disruption of the cellular gradients (e.g., the proton motive force) and unwanted exchange of molecules across the membrane. To assess the importance of T3SS pore gating for bacterial physiology, we therefore compared the fitness of several mutants of the FliP M-loop expressed from their native chromosomal location. Mutations that caused leakage through the pore induced a growth defect (Figure 7B), highlighting that despite the low number of secretion pores per cell, absence of gating caused a greatly reduced fitness.

**Figure 7.**
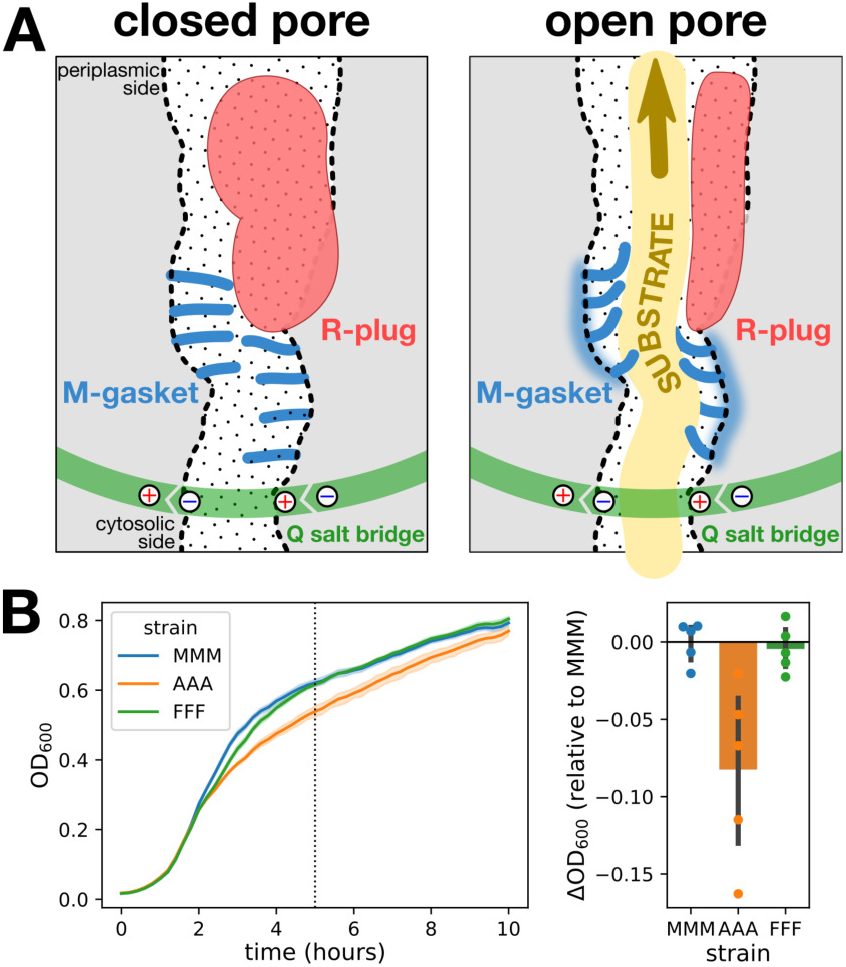
The T3SS secretion pore conserves membrane barrier during type-III secretion to maintain fitness. **A.** Model of T3SS secretion pore regulation to prevent membrane. In closed conformation, the M-gasket (blue) and the R-plug (red) cooperate to seal the channel and prevent alteration of membrane barrier. Upon substrate translocation, the M-gasket and R-plug undergoes conformational changes to enable opening the channel. Salt bridges in FliQ are critical to support M-gasket deformation and maintain the secretion pore tightly assembled. **B.** The loss of membrane barrier caused from leaky T3SS secretion pores results in altered fitness. Left: growth of several M-loop variants was followed over time by monitoring the OD_600_. The arrow indicates the growth phase (OD_600_ = 0.3) at which expression of the flagellar master regulator FlhDC starts increasing (18). Right: the AAA mutant, that is impaired in secretion pore gating, has a growth defect compared to the WT (MMM) and the FFF variants. Experiments were performed with at least 3 biological replicates. Error bars represent the standard deviation.

## Discussion

The type-III protein secretion machinery allows for the assembly of megadalton nanomachines such as the flagellum and related injectisome in the bacterial cell envelope. The assembly process of these complex nanomachines requires the secretion of dozens of protein subunits of various sizes and stoichiometries at a very high speed. Importantly, the T3SSs of the flagellum and the injectisome are capable of translocating substrate proteins across the inner membrane with a remarkable speed of several thousand amino acids per second, which is several orders of magnitude faster than protein export in other pore-based protein channels (1, 19). Furthermore, engineering of this secretion system has demonstrated that a wide range of proteins can be transported as substrates (20). Accordingly, it appeared reasonable to assume that a complex multi-level gating mechanism has evolved to maintain membrane integrity and to prevent the leakage of water and small molecules during high-speed substrate protein secretion.

The structures of the closed secretion pores of the T3SSs of the flagellum and the injectisome suggested that a plug domain in FliR and a stretch of methionine residues in FliP form constriction points that might contribute to gating of the T3SS (8). However, functional evidence of how the T3SS is able to facilitate the high-speed transport of substrate proteins while preventing the leakage of small molecules was lacking. In particular, the contribution of the highly conserved stretch of methionines in FliP in maintenance of the membrane barrier during active secretion remained unclear.

In this work, we performed an extensive functional analysis of the structural features of the T3SS secretion pore that were previously hypothesised to contribute to the gating mechanism. Our results demonstrate that the M-gasket made of five repetitions of a conserved loop of methionines in FliP (M-loop) and the R-plug domain in FliR are critical features of the T3SS secretion pore to preserve the membrane integrity during high-speed protein translocation. As suggested by the previous structures of the closed secretion pore, both the M-gasket and the R-plug seal the channel in its closed conformation, which renders any protein translocation impossible. Thus, conformational changes of both domains are required to enable substrate export. The FliR plug appears to be dedicated to the gating function, and its deletion does not otherwise affect flagellar function. In contrast, alterations in the M-loop frequently cause functional defects, supporting the idea that the M-gasket plays a dual role during T3SS function. It acts both as a flexible gasket to maintain the membrane barrier during substrate protein translocation, and as a hinge to accommodate pore opening and substrate translocation. The convoluted spatial organization of the M-gasket in the channel attributes a unique position to each of the 15 methionine residues of the M-loops, that is spread over 20Å in the secretion channel. This suggests that different residues in each M-loop might contribute to the formation of the M-gasket over the different events of pore opening and substrate translocation.

Using extensive targeted and random mutagenesis of the M-loop, we demonstrated that a complex combination of physicochemical features of residues forming the mouth of the T3SS secretion pore is required to support membrane gating and substrate secretion, and that unlike any other amino acid, methionines ideally fulfil this requirement. Moderate hydrophobicity, a long side chain, and side chain flexibility are likely important to maintain a dynamic interaction between the FliP subunits and to prevent leakage by maintaining a close contact with the secreted substrates while accommodating conformational changes required for their rapid translocation. This immediate proximity with the substrates is likely a reason why a net charge is not tolerated in the channel.

Maintaining membrane barrier during protein secretion is crucial to bacterial fitness. This was shown for the bacterial Sec translocon that involves hydrophobic residues and a plug domain to gate the constricted section of the channel (3). We report here a similar observation for the T3SS in which a leaky M-gasket induces a growth defect. This is further highlighted by the fact that only a handful of T3SS secretion pores are assembled in the cell envelope. Interestingly, while active secretion reduced leakiness in mutants of the Sec translocon, we observed the opposite in the fT3SS. Leakage caused by a mutation in the M-loop of FliP occurred only in actively secreting fT3SS.

The close structural homology between the three components of the secretion pore makes it tempting to hypothesize that the T3SS secretion pore evolved sequentially from a simpler pore, by specialization of the FliR subunit and of its plug domain. However, the potential function and impact of specific adaptations on secretion pore gating, such as the extra methionine residues in the M-loop of some virulent T3SS homologues, or conversely the existence of rare flagellar homologues with reduced number of methionines in the M-loop remains to be investigated.

## Material & Methods

### Bacteria, plasmids and media

Strains used in this study derived from *Salmonella enterica* serovar Typhimurium strain LT2 and are listed in Supplementary Table S3. Lysogeny broth (LB) contained 10 g of Bacto-Tryptone (Difco), 5 g of yeast extract, 5 g of NaCl and 0.2 ml of 5N NaOH per litre. Soft agar plates used for motility assays contained 10 g of Bacto-Tryptone, 5 g of NaCl, 3.5 g of Bacto-Agar (Difco) and 0.2 ml of 5N NaOH per liter.

### Chromosomal modifications

Chromosomal mutations were obtained using lambda-RED mediated recombineering (21) and a Kan-SceI cassette as selection marker (22). Site-specific chromosomal insertion of the Kan-SceI cassette and subsequently the replacement with the desired chromosomal mutation was achieved by inducing the expression of the λ-Red phage genes (gam, bet, exo, encoded on the temperature-sensitive plasmid pWRG730), by heat-shock. Both, the Kan-SceI cassette and the replacement fragments were amplified by PCR with flanking homologous regions of 40 bp to the target region. The desired chromosomal modification was obtained by counter-selection on LB plates containing anhydrotetracycline to induce expression of the meganuclease I-SceI. Recombinant colonies were verified by PCR, followed by sequencing.

### Random mutagenesis

Random mutagenesis of the FliP M-loop was achieved by using a degenerate oligonucleotide with a homologous overhang complementary to the sequence immediately upstream of the M-loop (cgacctggtgatcgccagcgtattgatggcgttgggg **NNS NNS NNS** gtgccgcca-gcgac). The **NNS** codon allows to obtain all possible amino acids with limited redundancy and avoiding rare and stop codons. Mutagenic fragments were amplified by PCR using the degenerate oligonucleotide together with a reverse primer (GCTGATAATGAGGCCGGTAA), and were subsequently recombined on the chromosome using the lambda-RED procedure. Selection for functional variants was carried out either by selecting for ampicillin-resistant clones in a genetic background containing a substrate fused to the β-lactamase (see. Quantification of substrate secretion) or by picking and subcloning flares of motile bacteria after short-term growth in soft agar. Alternatively, whether variants were able to assemble a functional flagellum was determined using a genetic reporter (see. Genetic reporter of complete flagellum assembly).

### Motility assay in soft agar

In order to assess the motility of the various *Salmonella* mutants, overnight cultures were inoculated in soft agar plates with a pin tool and incubated at 30°C. Plates were scanned automatically from within the incubator at regular intervals using the light transmission mode of a Perfection V800 Photo scanner (Epson) controlled by a custom software. Motility was determined from the radius of the rings of cells originating from the points of inoculation. Briefly, background was subtracted using an image of the plate shortly after inoculation as a reference. Motility rings were detected using ImageJ by applying a threshold (Huang algorithm) and automatic detection of the contours of the rings.

### Quantification of substrate secretion

Mutations of interest were transduced in a strain carrying a Δ*flgBC* mutation and a translational fusion of the hook protein FlgE to the β-lactamase TEM-1 lacking its secretion signal (FlgE-bla) (23). This permitted secretion of FlgE-bla directly into the periplasm and conferred resistance to ampicillin proportionally to the amount of secreted substrate. The ampicillin IC50 value was determined by growing the bacteria in a 96 wells plate with a titration of ampicillin (0 to 1024 μg/mL) and by fitting a logistic function on the endpoint OD_600_.

### Kinetics of water movement across the inner membrane

Bacteria were grown in LB medium containing 10 g/L NaCl until reaching an OD_600_ of 0.6. Cells were washed twice in 1×PBS, concentrated to 30 OD_600_/mL, and diluted 1:1 in a high concentration of salt (*e.g.*, 1M guanidinium in 1×PBS). Absorbance at 600 nm was monitored immediately using using Cytation3 or Synergy H1 plate readers (Biotek), and variation over time reflected the flux of water across the inner membrane. The initial speed of efflux was quantified, after standardization to the OD unit, as the slope of the initial tangent to the curve (ΔAbs/min).

### Genetic reporter of complete flagellum assembly

Complete assembly of the flagellum was assessed by reporting the genetic switch to flagellar class III genes. For this, a transcriptional fusion of the FljB flagellin and the mudA transposon, that carries the β-galactosidase (LacZ) gene (24), was used. Bacteria able to assemble the flagellar hook and switch to late substrate expression thus appeared as red colonies on MacConkey agar plates (Difco).

### Immunostaining of flagellar filaments

To determine the number of flagella on individual bacterial cells, flagellar filaments were labeled with anti-flagellin antibodies coupled to organic fluorochromes. Briefly, bacteria were grown till mid-log phase and flagellar filaments were incubated with polyclonal anti-FliC (ref. 224741) and anti-FljB (ref. 228241) antibodies (Becton Dickinson & Company). Subsequently, the secondary antibody α-rabbit conjugated with Alexa Fluor 488 (Invitrogen) was added. Bacterial DNA was stained with Fluoroshield containing DAPI (4’,6-diamidino-2-phenylindole; Sigma). Filaments were imaged using a Zeiss Axio Observer Z1 inverted microscope at 100× magnification with an Axiocam 506 mono CCD-camera and the Zen 2.6 pro software. The images were further processed with ImageJ.

### Bacterial fitness

Bacterial fitness was determined from the difference in growth of mutant strains compared to the WT reference. Growth was monitored by measuring every 5 minutes the optical density of liquid cultures cultivated in 96 well plates at a wavelength of 600 nm using Cytation3 or Synergy H1 plate readers (Biotek).

## ACKNOWLEDGEMENTS

We would like to thank the Erhardt and Charpentier laboratories for continuous help and support, useful discussions, and for critical comments on the manuscript; Tim Sullivan and Heidi Landmesser for expert technical assistance; Tohru Minamino for providing us with the anti-FlhA_C_ antibody. This work was supported by the Deutsche Forschungsgemeinschaft (DFG) research grant no. ER 778/2-1 (to M.E.), and a fellowship from the Alexander von Humboldt foundation (to T.T.R.). The funders had no role in study design, data collection and analysis, decision to publish, or preparation of the manuscript.

## AUTHOR CONTRIBUTIONS

T.T.R. and M.E. conceived the project, designed the study, wrote and revised the paper. T.T.R., S.H., U.v.L., A.G., and D.F.B. performed the experiments. T.T.R., M.E., and D.F.B. analyzed and interpreted the data with input from all authors. E.J.C.G. contributed to the bioinformatic analyses. E.C., M.E. and T.T.R. contributed funding and resources. All authors commented and approved the manuscript.

## Supplementary Information

**Figure S1.**
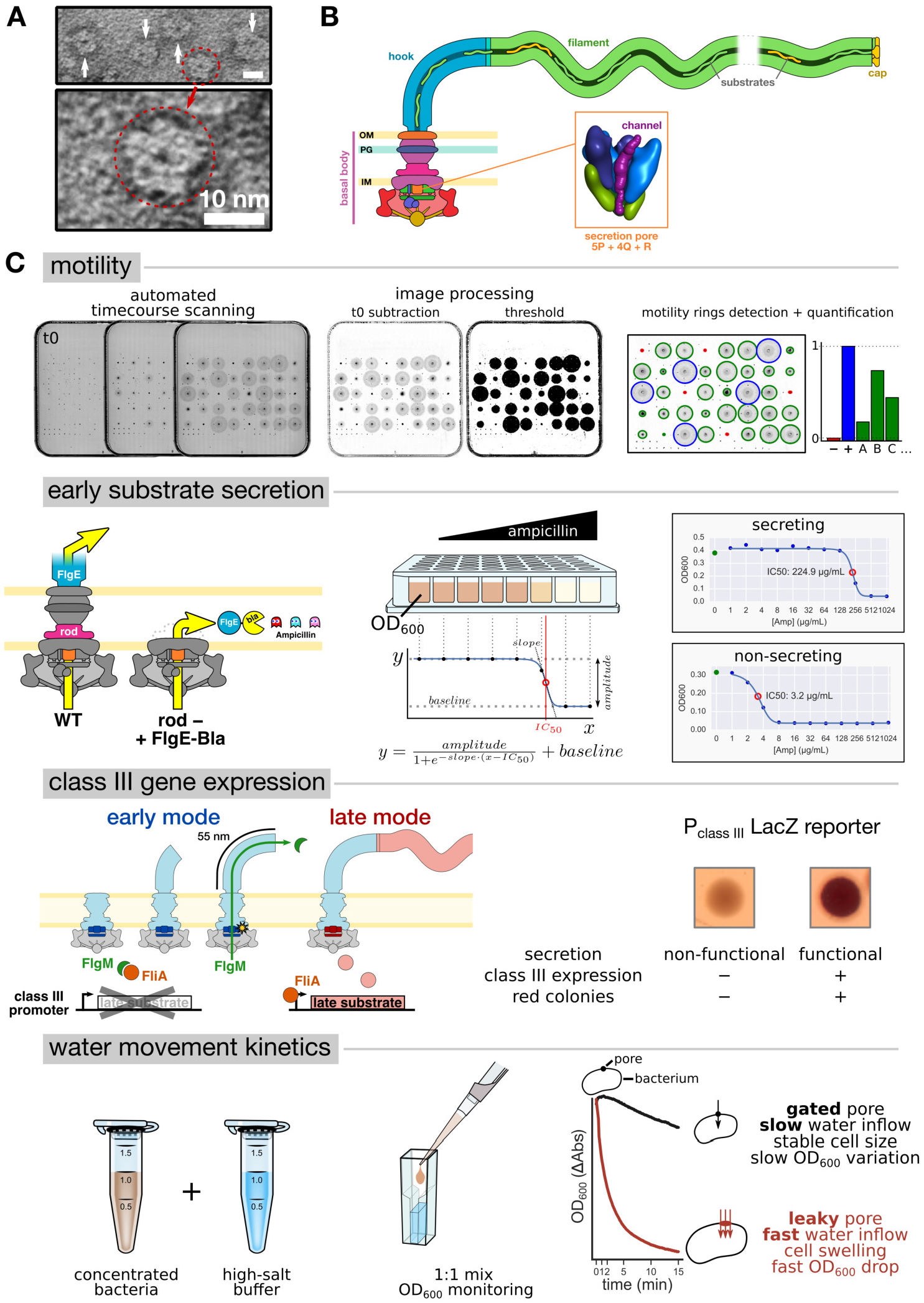
**A.** Over-expressed FliP form oligomers reminiscent of a pore. **B.** Location of the T3SS secretion pore in the assembled flagellum. The FliP1, FliQ1, and FliR subunits are hidden to better visualize the inside of the pore. The channel (shown in purple) was computed using Caver 3 (see Figure 2). **C.** Schematic description of the assays used in this study. Motility: bacteria are inoculated in soft agar and plates are monitored for the apparition of motility rings. Scanned images of the motility plates are processed to subtract background and detect the contour of the motility rings. Quantification is done by measuring the average radius of the motility rings and using ΔfliP and WT FliP as negative and positive controls, respectively. Early substrate secretion: Deletion of the rod proteins FlgB/C enables secretion of the flagellar substrates in the periplasmic space. Fusion of the β-lactamase TEM-1 (devoid of its secretion signal) to an early flagellar substrate (the hook protein FlgE), renders the cells resistant to ampicillin to a level that is proportional to the secretion of the fusion substrate. The level of ampicillin resistance is quantified by monitoring bacterial growth in presence of increasing concentrations of ampicillin, and by fitting the observed data to a logistic function. The fitted IC50 parameter represents the antibiotic concentration that inhibits bacterial growth by 50%. Class III gene expression reporter: Flagellar assembly occurs in two consecutive steps. First the basal body and hook (early substrates) are assembled, and during this time expression of the late substrates is repressed by the anti-sigma factor FlgM. Upon completion of the hook, FlgM is secreted and relieves the inhibition of flagellar class III promoters, that control the expression of late substrate, to enable assembly of the flagellar filament. Placing the LacZ gene under the control of a class III flagellar promoter allows to report the successful assembly of the hook-basal body, and thus the proper function of the T3SS secretion pore. Functional clones appear red on MacConkey plates. Water movement kinetics (secretion pore leakage): Bacteria expressing different variants of the T3SS pore are washed and concentrated in 1×PBS. The bacteria are then diluted 1:1 in a high salt buffer. Variants bearing a functional, sealed, T3SS pore are resistant to passage of water across the pore and their cell size thus remains stable. Conversely, variants with a defect of pore gating enable a flow of water across the pore, which results in cell swelling and a rapid drop of optical density.

**Figure S2.**
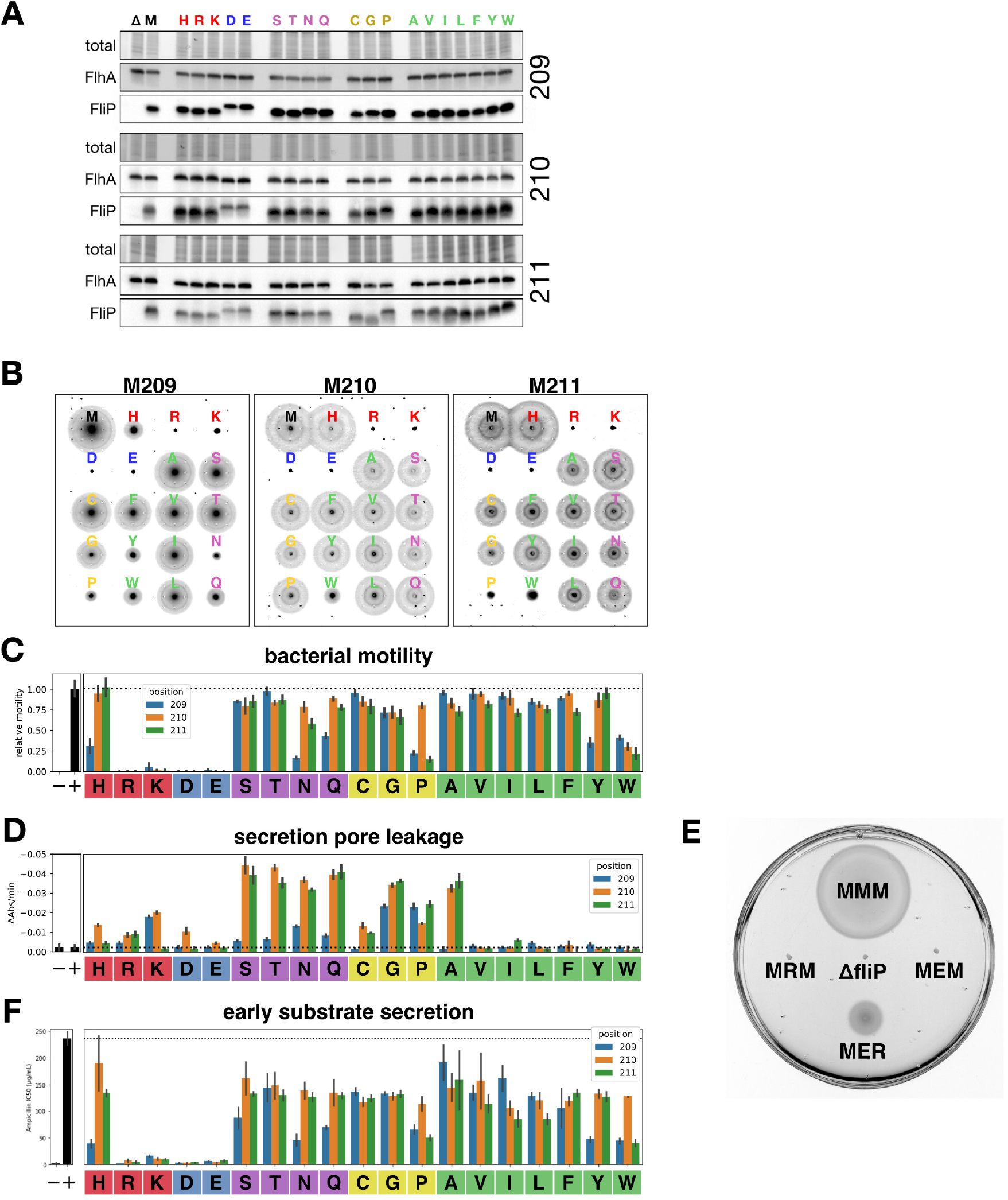
**A.** All single amino-acids substitutions in FliP MMM (209–211) are expressed and localized to the membrane. Western-blot of membrane extracts. A 3×FLAG tag and flexible linkers (SAGASA-DYKDHDGDYKDHDIDYKDDDDK-SAGASA) were inserted in a non-conserved region (after G157) to allow detection of the protein. Membrane fractions were prepared by mechanical disruption of the cells and differential centrifugations. TCE staining of total proteins and detection of the membrane flagellar protein FlhA were used as loading controls. Furthermore, FlhA assembly requires the prior correct assembly of FliP (1). Δ: ΔfliP strain; M: W.T strain (MMM) **B.** Motility in soft agar of all single variants of the M-loop. Scanning of the plates was automated and background was subtracted from the initial time point. **C—D.** Aggregated data of the quantification of several replicates of the motility (**C**) and leakage (**D**) assays as described in Supplementary Figure 1C. Assembly of the flagellum and maintenance of membrane barrier are two crucial functions of the T3SS secretion pore that are affected by the presence of certain residues in the M-loop. **E.** Motility in soft agar of two non-motile charged variants of the M-loop (M210E and M210R), and of a MER revertant that derives from the M210E background. The WT (MMM) and a strain lacking FliP are shown as positive and negative controls, respectively. **F.** Secretion in the periplasm of a fusion between the hook protein (FlgE) and the β-lactamase TEM-1 devoid of its signal peptide renders the strain resistant to ampicillin.

**Figure S3.**
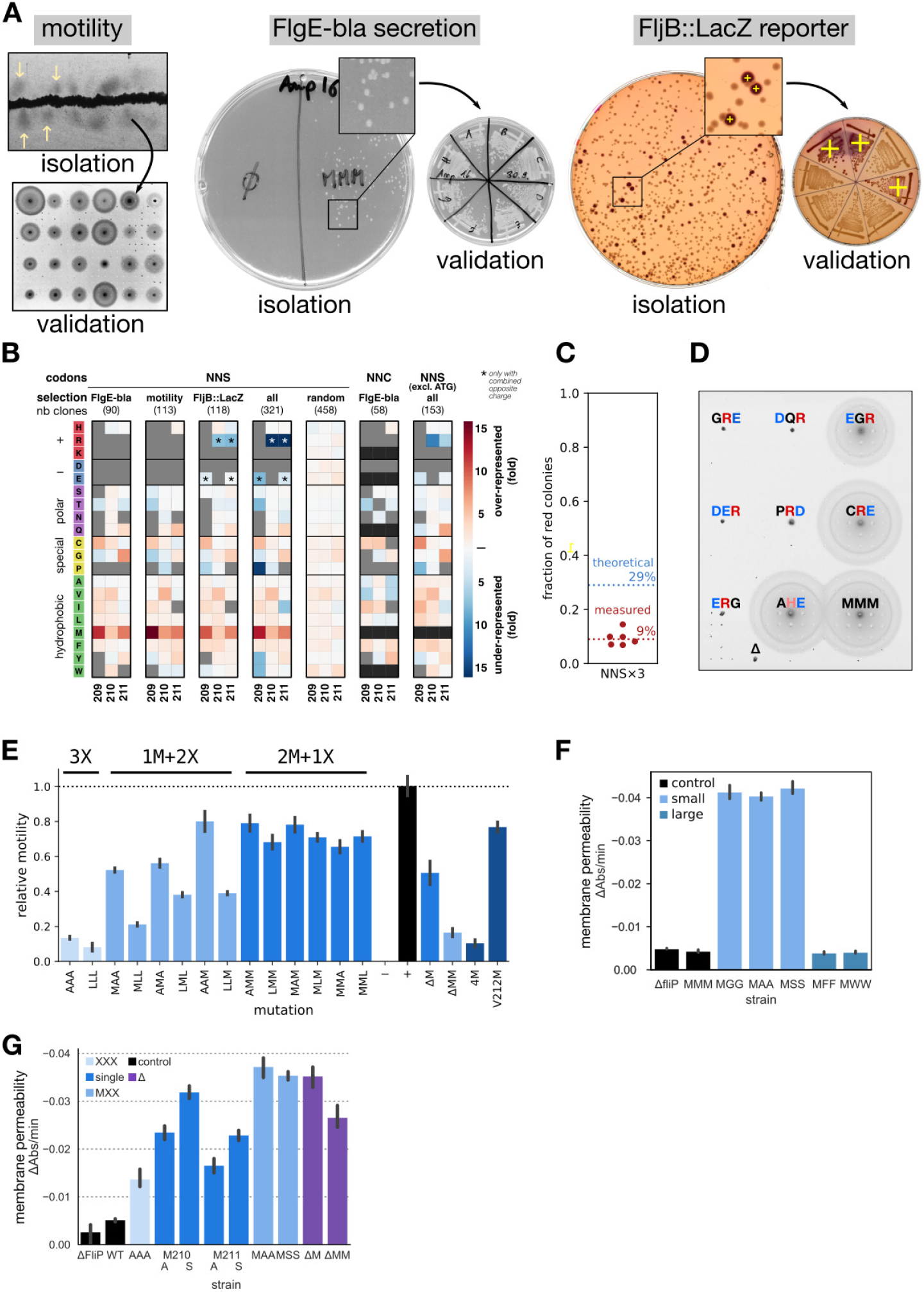
**A.** Independent methods of functional random M-loop variants selection. All methods are performed after recombination of a degenerated NNS×3 oligonucleotide in place of the M-loop. **Left:** motility selection relies on the successful assembly of the bacterial flagellum, which confers the ability to swim away from the inoculation site. **Middle:** FlgE-bla secretion confers ampicillin resistance only when the T3SS is able to secrete the fusion substrate. **Right:** In this reporter system, clones are first selected for successful recombination. Functional clones are identified using a MudA translational fusion to the flagellin FljB, that triggers β-lactamase expression upon completion of the flagellar hook-basal body. Positive clones appear red on MacConkey agar. **B.** Frequency of amino-acid use per position in the random M-loop variants. From left to right: FlgE-bla/motility/FljB::LacZ: three independent methods of selection for functional motifs. All: combination of the result of the three selection methods. Random: absence of selection. Those last two datasets indicate that in absence of methionine a slight over-representation in medium size hydrophobic amino acids (all positions), cysteine at position 209, and glutamine/glycine (position 211) is observed, without reaching the level of methionine. NNC/FlgE-bla: FlgE-bla based selection using an NNC×3 degenerated oligonucleotide to avoid methionines. NNS excl. ATG: exclusion of the motifs that do not contain methionine. **C.** The FljB::MudA reporter system enables to measure the proportion of functional M-loop variants by counting the proportion of red colonies. Using an NNS×3 recombination oligonucleotide and excluding amino-acids not tolerated at each position (209: HRKDEWYPNQ (13 codons), 210: RKDEW (7 codons), 211: RKDEWP (9 codons), see. Figure 2B) plus one stop codon, a functional motif would be obtained with a probability 18/32×24/32×22/32 = 0.29, which is much higher than the observed frequency. This suggests that not only the nature of the residues at each position, but also their combination has an effect on the function of the M-loop. **D.** Example of M-loop motifs with a combination of negative and positive charge. Depending on the position and residues, the phenotype can be non-motile (GRE, DQR, DER, PRD, ERG) or motile (EGR, CRE, AHE), which confirms that a net charge in the M-loop does not allow T3SS function. **E.** Combination of methionines and leucines or alanine indicate that T3SS function is roughly proportional to the number of methionine. ΔM/ΔMM: deletion of 1 or 2 methionines in the M-loop. 4M: insertion of an extra methionine in the loop. V212M: substitution of the valine 212 that is immediately after the M-loop to mimic the MMMM motif from the SpaP protein in the vT3SS-1 of *S.* typhimurium. **F.** Among the MXX M-loop mutants, only variants to residues with small side chain induce a leakage through the secretion pore. **G.** Single mutations to serines induce a stronger leakage than their alanine counterparts, indicating that leakage is favored by hydrophilicity and hindered by hydrophobicity, as expected. Unlike triple mutations, double mutations of only the two exposed methionines reached a maximum of leakage similar to that observed in our previously reported single methionine deletion mutant (ΔM). A double deletion mutant (ΔMM) was however less leaky, likely resulting from a structural effect rendering the complex unstable as observed for the AAA mutation.

**Figure S4.**
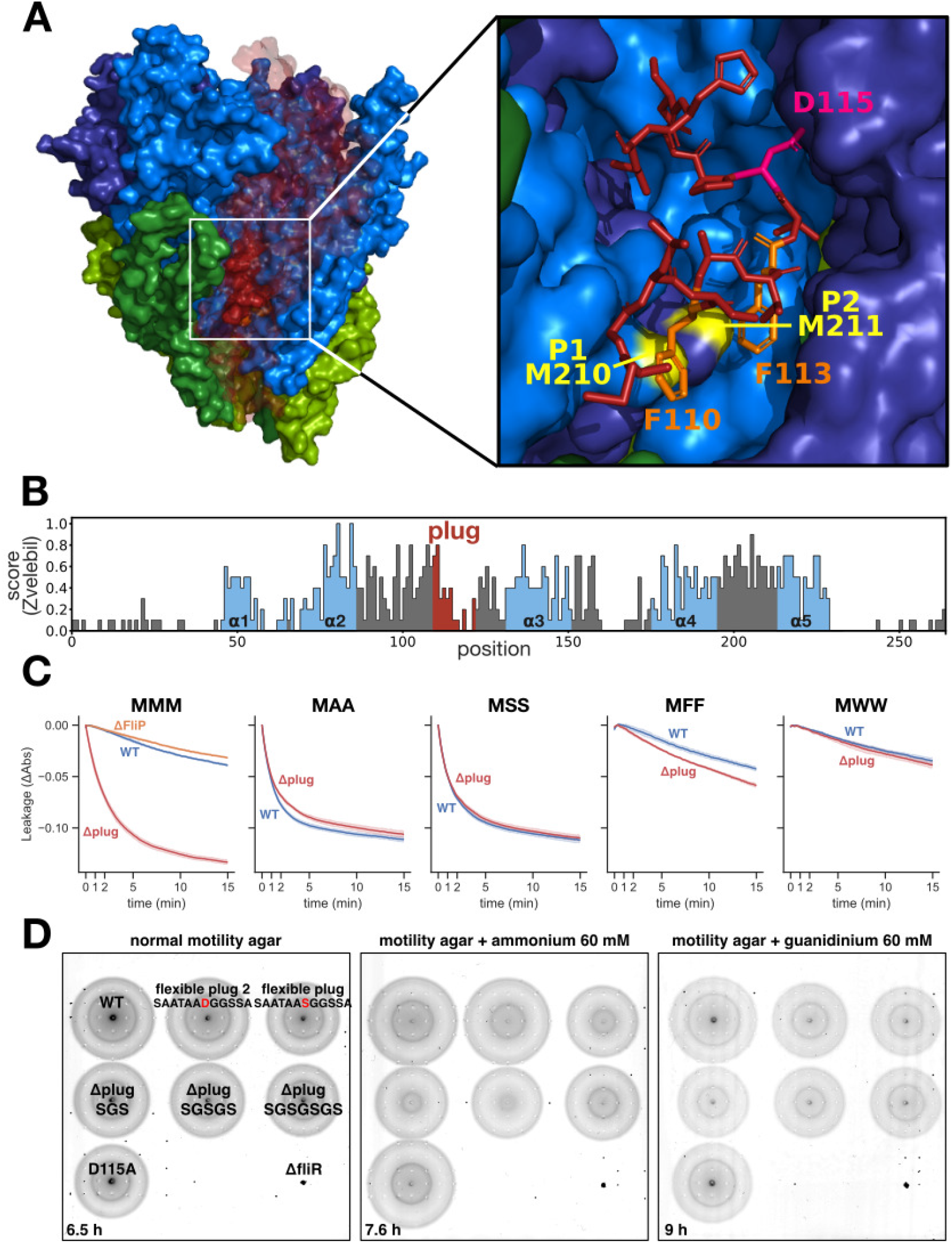
**A.** The plug domain of FliR is in direct interaction with the M-gasket. In particular, two mildly conserved hydrophobic residues (F110 and F113) are in contact with two methionines (M210 of P1 subunit and M211 of P2 subunit). Another residue of the plug D115, that is highly conserved, is also in interaction with T80 in the P4 subunit. Only the plug residues of FliR are represented for clarity. **B.** The plug domain of FliR displays a low Zvelebil amino-acid conservation score, which reflects a high diversity of amino-acid properties among homologs (2). **C.** The combination of the plug of FliR and the M-gasket is required to maintain membrane barrier. Failure of either of the domains is sufficient to enable leakage of small molecules through the secretion pore. **D.** Motility in soft agar, with or without addition of 60 mM ammonium/guanidinium chloride in the medium, reports that various mutations of the FliR plug do not result in a marked alteration of flagellum assembly and function.

**Figure S5.**
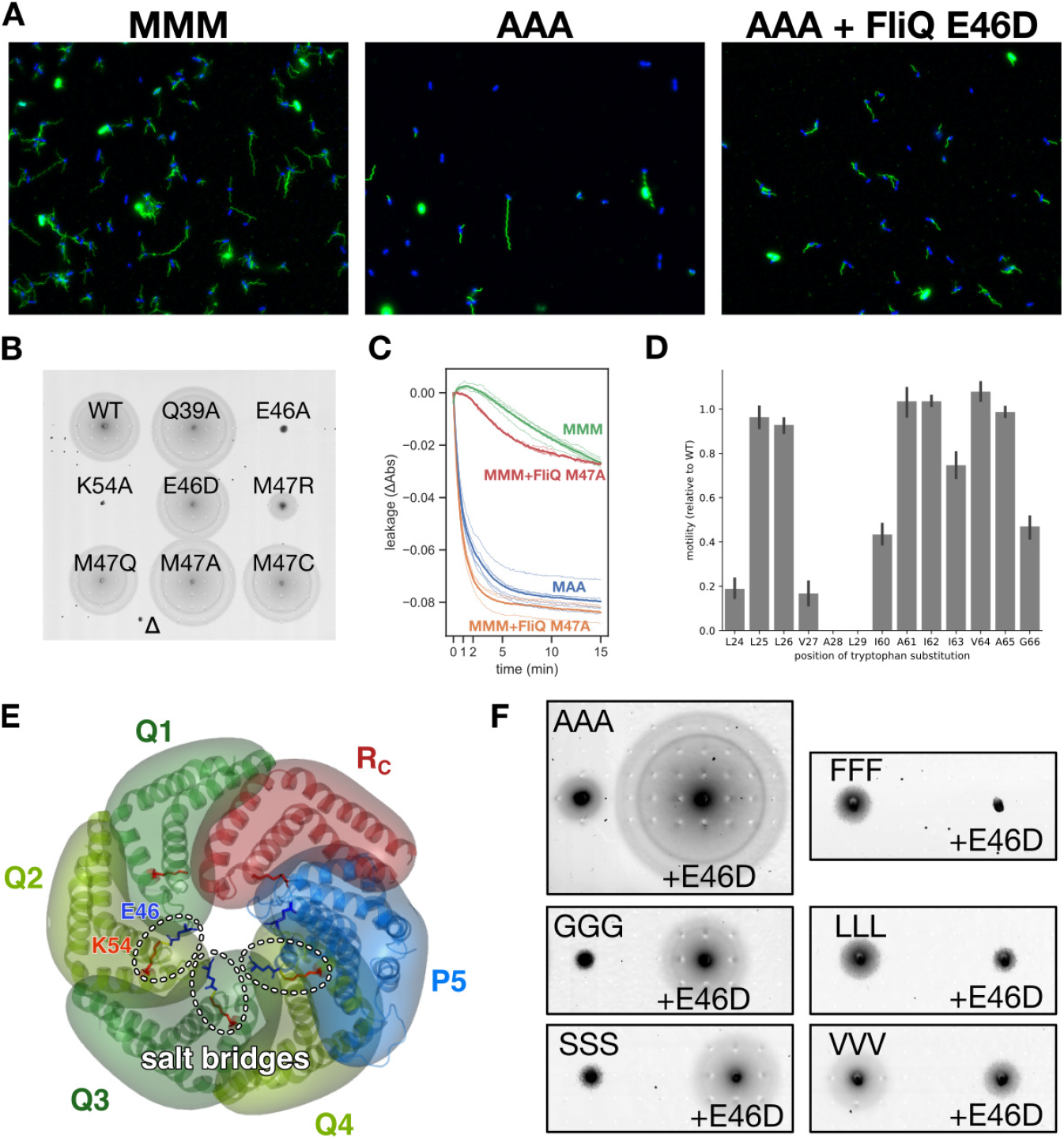
**A.** Representative images of flagellum immunostaining (green color). The AAA M-loop mutant has a structural defect and can assemble less filaments than the WT (MMM), thus resulting in altered motility. The bacteria are stained with DAPI (blue). **B.** The E46-K54 residues involved in an inter-subunit salt bridge are required for T3SS secretion pore function. Conversely, the Q39 and M47 residues do not play a crucial role. **C.** The M47 of FliQ does not play an important role in fT3SS secretion pore gating. **D.** Tryptophan scanning of the central parts of FliQ α-helices. The A28W and L29W mutations abrogate motility. **E.** The three FliQ inter-molecular salt bridges are located close to the mouth of the channel and participate in maintaining the stability of the M-gasket. **F.** The FliQ E46D mutation improves the function of the AAA/GGG/SSS M-loop variants and conversely further impairs motility of the FFF/LLL/VVV variants.

**Table S1.**
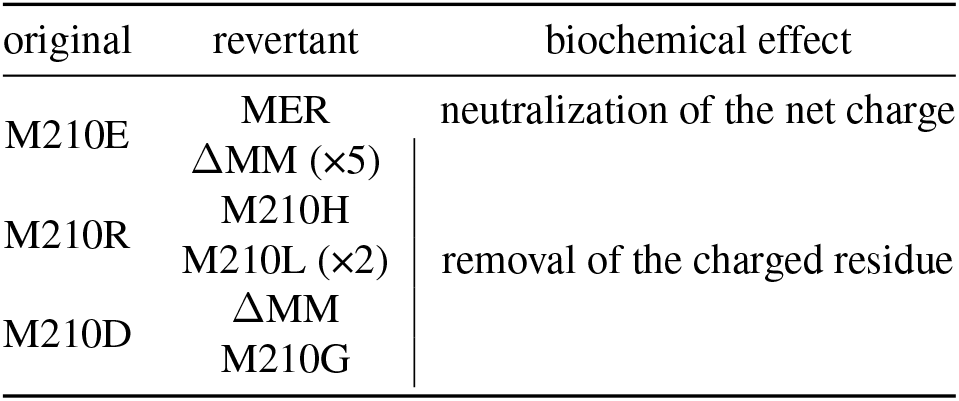
Revertants of M-loop variants with charged residues

**Table S2.**
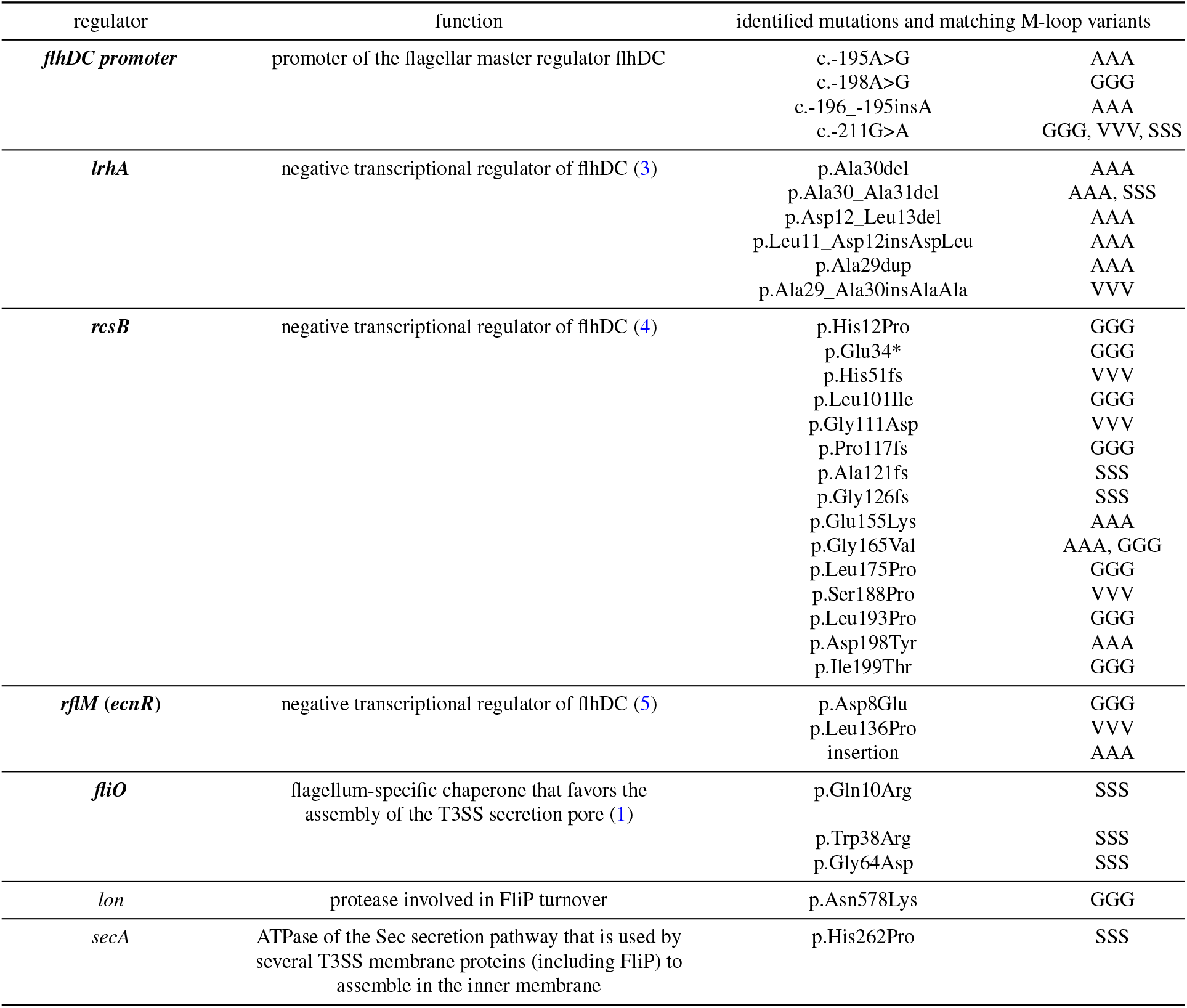
List of mutations in known regulators of flagellum assembly that enhance function of impaired FliP M-loop variants. Regulators that were identified independently in several backgrounds or experiments are displayed in bold. The [HGVS nomenclature](https://varnomen.hgvs.org/) is used.

**Table S3.**
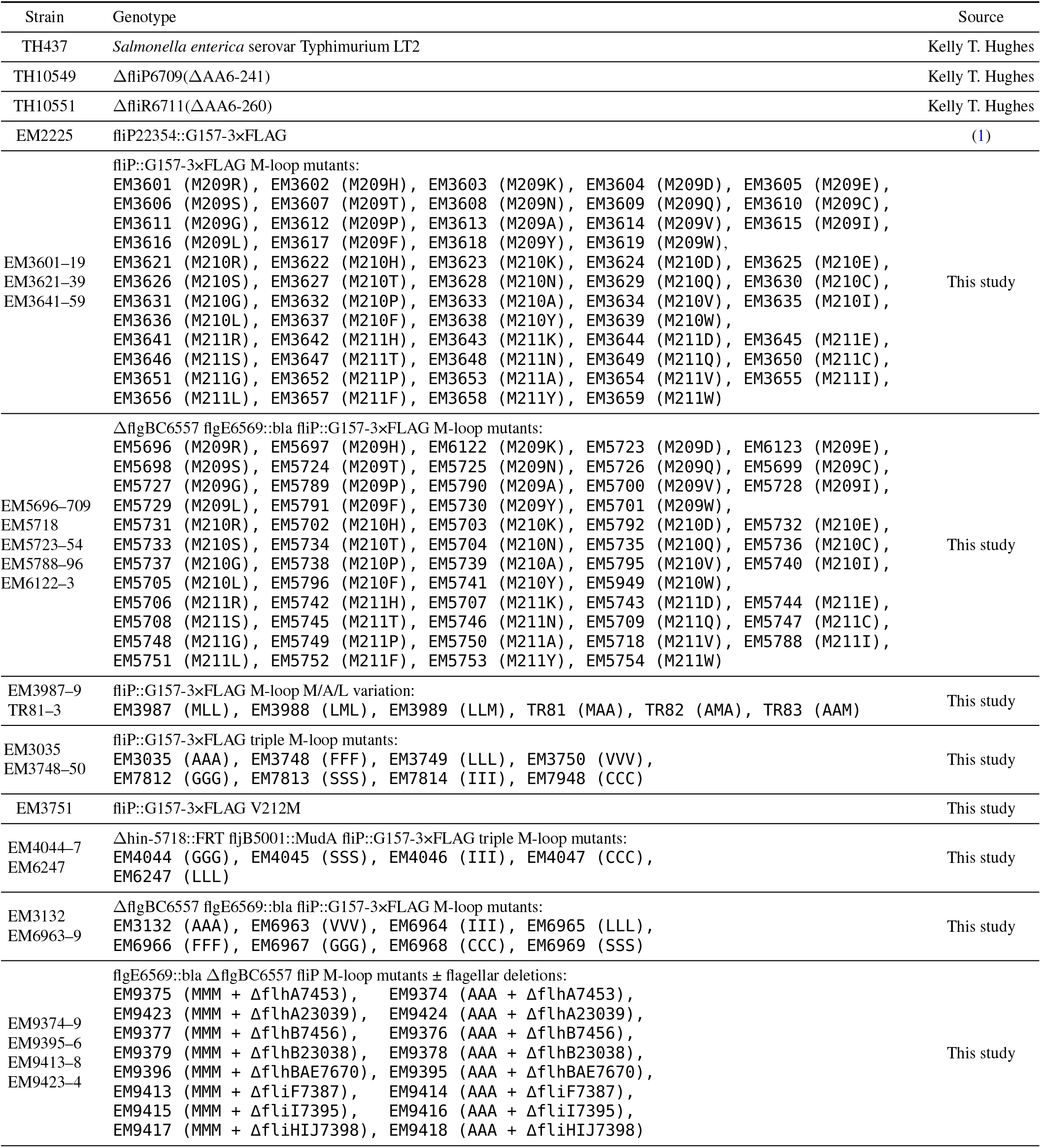
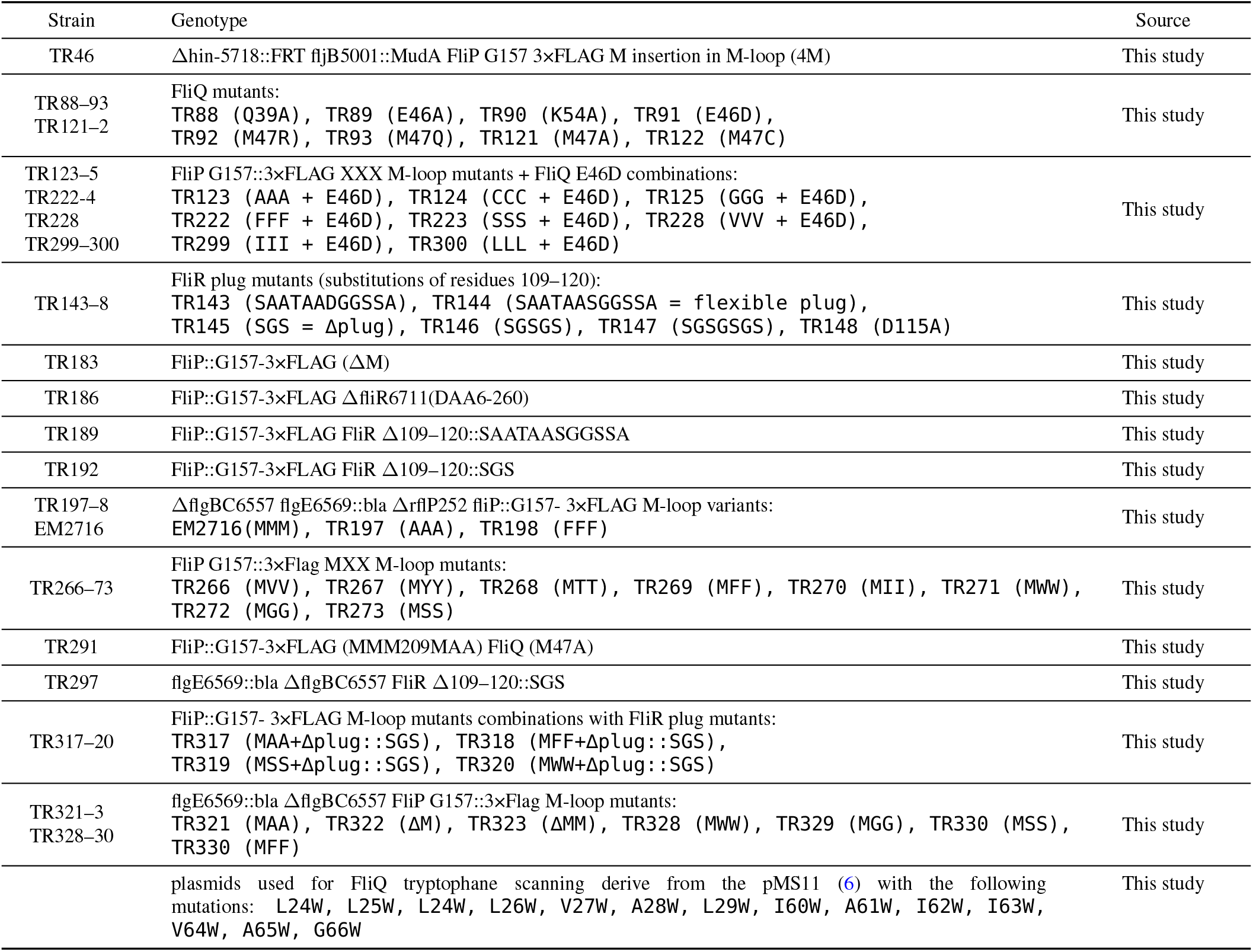
List of strains used in this study (without random mutants and revertants)

